# *Mycobacterium tuberculosis* partitions the Krebs cycle under iron starvation

**DOI:** 10.1101/2025.05.12.653400

**Authors:** Agnese Serafini, Acely Garza-Garcia, Davide Sorze, Luiz Pedro Sorio de Carvalho, Riccardo Manganelli

## Abstract

In this study, we investigated how iron limitation alters central metabolism in *Mycobacterium tuberculosis* using metabolomics and stable isotope tracing. Our findings reveal a well-orchestrated metabolic program to enable Krebs cycle activity despite the inefficient action of its iron-dependent enzymes. Under such conditions, carbon flux through the oxidative branch of the Krebs cycle is stalled, resulting in the accumulation of metabolites that are partially secreted. As a result, carbon flux from glycolysis is partially diverted to the reductive branch of the Krebs cycle to support the production of oxaloacetate and malate through the activity of phosphoenolpyruvate carboxykinase and pyruvate carboxylase. Both branches terminate with the synthesis of malate, which is secreted. This unprecedented split of the Krebs cycle and malate secretion in a bacterial pathogen facilitates the continuous flow of carbon through the core of carbon metabolism, overcoming the metabolic stalling triggered by iron starvation.

---

Over the past decade, mounting evidence has indicated a clear correlation between central metabolism and antibiotic resistance. It has been demonstrated that clinically relevant mutations in genes encoding core metabolic enzymes are associated to antibiotic resistance (Lopatkin et al. 2021) and tolerance (Hicks et al. 2018). Furthermore, secondary or “collateral” effects of antibiotic action involve alterations to central carbon metabolism (CCM) (Wang et al. 2019; Kawai et al. 2019) and specific metabolic states affect their efficacy (Allison et al. 2011; Lim et al. 2021; Lopatkin et al. 2019; Schrader et al. 2021).

*Mycobacterium tuberculosis* (*Mtb*), the causative agent of tuberculosis (TB), has been recently included in the Bacterial Priority Pathogen List of the World Health Organization (WHO 2024) due to its intrinsic resistance to antibiotics and the increased spreading of multi-drug-resistant strains. Since the 1960s, only three new drugs have been approved for the treatment of TB (Murray et al. 2015; Zheng & Av-Gay 2016; Dartois & Dick 2024), and resistance to these new antibiotics is increasing. It is therefore critical that new medicines are developed to treat this infection. For more efficient antibiotic discovery, we must gain a more comprehensive understanding of the physiology and metabolism of *Mtb*. This knowledge will help identify novel vulnerabilities and inform the design of more effective therapeutic strategies.

During infection pathogenic bacteria are subject to nutritional immunity, a process in which the host alters metal availability to intoxicate or starve the invading pathogens (Murdoch & Skaar 2022). Metals are essential micronutrients that play structural, signalling or catalytic roles in most of cellular processes. Many enzymes of central metabolism depend on metals for their activity, however the effects of nutritional immunity on CCM have not been extensively investigated (Serafini 2021). Experimental evidence indicates that *Mtb* is exposed to iron starvation during infection (Kurthkoti et al. 2017; Wells et al. 2013; Fang et al. 2015; Madigan et al. 2015; Pisu et al. 2020). This study aims to examine the relationship between iron homeostasis and CCM, a complex and understudied aspect of *Mtb* metabolism. *Mtb* exposed to prolonged and severe iron starvation exhibits a reduction in growth rate until replication is arrested, yet, cells remain viable (Kurthkoti et al. 2017). Based on the publicly available transcriptomic and proteomic data of H37Rv *Mtb* strain grown under iron limitation (**Dataset S1** and Kurthkoti et al. 2017; Wong et al. 1999; Serafini et al. 2013), we have recently proposed a model that explains how this pathogen might adapt its core of CCM to limited iron availability (Serafini 2021). We proposed that *Mtb*: i) reduces the activity of the Krebs cycle due to the compromised activity of its iron-dependent enzymes, which consequently reduces the production of FADH_2_, NADH and ATP; ii) activates an iron-independent pathway, the glyoxylate shunt, to maintain succinate and malate production; iii) secretes succinate to maintain an energised cell membrane, bypassing the reduced ATP synthase activity.

The aim of the present study is to test these predictions thereby clarifying the effect of iron starvation on *Mtb* metabolism. We monitored metabolite levels using liquid chromatography coupled to mass spectrometry and employed ^13^C isotope tracing to identify the metabolic pathways active under iron starvation. These experiments were conducted under the same growth conditions used in the more comprehensive transcriptomic study (Kurthkoti et al. 2017), which, like the other published studies on iron limitation (Wong et al. 1999; Serafini et al. 2013), included two carbon sources in the growth medium, glucose and glycerol. To circumvent genomic background-dependent effects, the work was performed in two *Mtb* strains with different metabolic capacities, H37Rv and Erdman (Serafini et al. 2019; Marrero et al. 2010).

## Results

### Severe growth defects accompany iron limitation

As demonstrated previously (Kurthkoti et al. 2017), *Mtb* exposed to severe iron starvation enters a non-replicative state that can persist for months. This severe iron starvation was achieved by subculturing bacteria in the absence of a source of Fe^3+^, followed by the addition of the Fe^3+^ chelator deferoxamine (DFO), to trap any residual ion. Equivalent conditions were used in our laboratory to reproduce the growth arrest phenotype in Erdman and H37Rv (**Figure 1A-D)**. As expected, the absence of a Fe^3+^ source (0 μM FeCl_3_) resulted in a growth slowdown, while the addition of DFO (0 μM FeCl_3_ + DFO) led to growth arrest (**Figure 1A and B**). In the latter, Erdman exhibited an apparent decline in viability over several weeks (**Figure 1D**) compared to H37Rv (**Figure 1C**). This decline could be attributed to a loss of viability, or more pronounced clumping of the cells in Erdman cultures (data not shown), that resulted in a reduction in the number of colony-forming units (CFUs), or to the cells entering an unculturable state (Kurthkoti et al. 2017).

**Figure 1.**
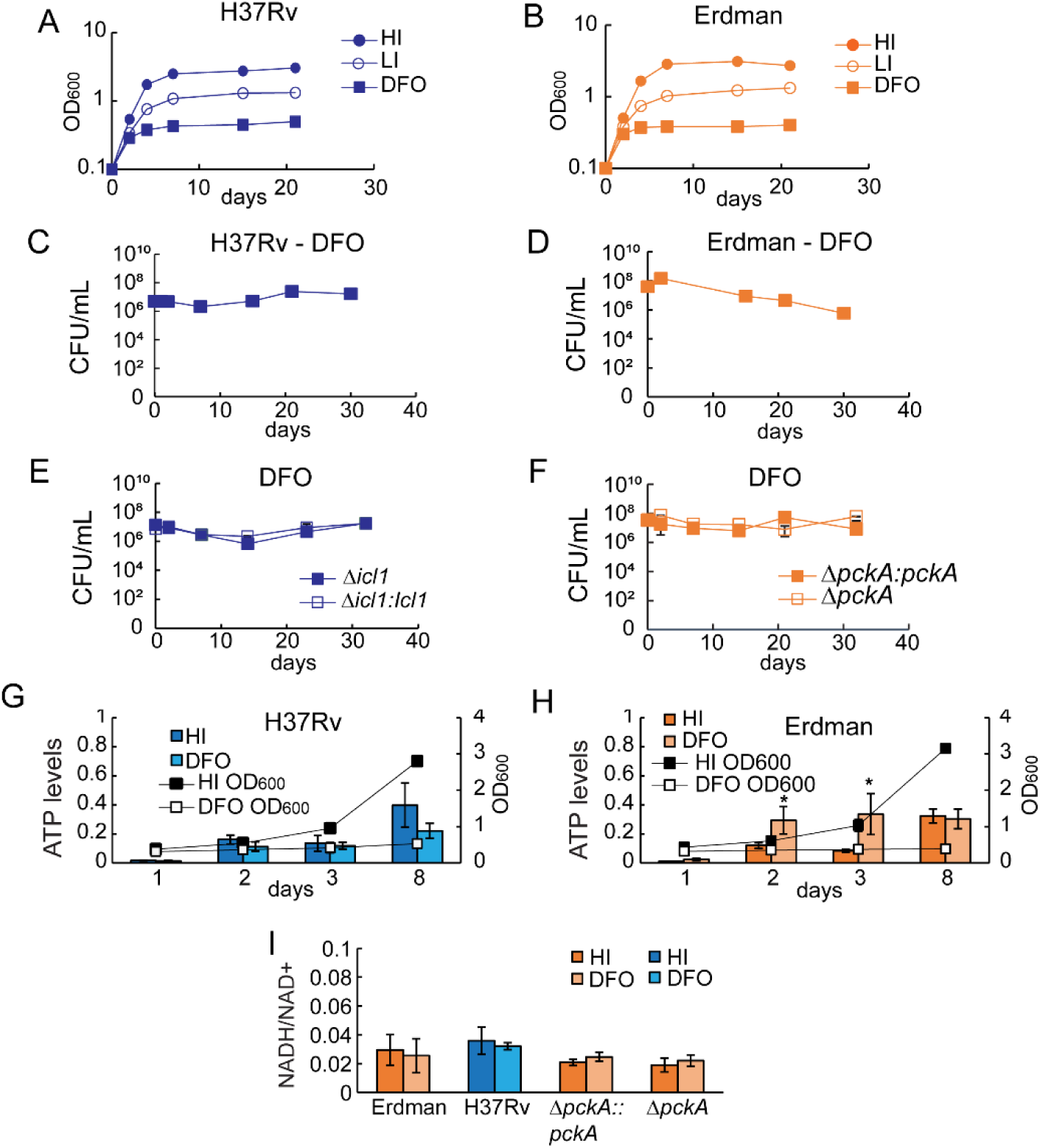
Survival to Fe^3+^ starvation in *Mtb* H37Rv and Erdman strains. Cells were exposed to 50 μM FeCl_3_ (HI: High Iron), 0 μM FeCl_3_ (LI: Low Iron,) or 0 μM FeCl_3_+ DFO (DFO). Growth was monitored for 3 weeks by measuring OD_600_ (A, B), and survival was monitored for more than 4 weeks by CFU/mL (C, D, E, F). The charts show one experiment representative of 2-3 independent experiments. The CFU/mL charts show average and average deviation of two technical replicates of one independent experiment. G, H) ATP levels and growth (OD_600_) after 1, 2, 3 and 8 days in DFO. ATP levels were calculated as µM of ATP molecules in about 10^7^ cells (0.1 optical density at 600 nm). The data are the average and standard deviation of three independent experiments and three technical replicates each. I) NADH/NAD+ ratio detected after 3 days of exposure to DFO. The data are the average and standard deviation of two independent experiments and two technical replicates. The p-values were calculated against the HI condition. * = p value < 0.05.

Previous studies have shown that replication-arrest-inducing stress conditions such as hypoxia and nutrient starvation (Eoh & Rhee 2013; Rao et al. 2008; Gengenbacher et al. 2010) reduce ATP production and alter the NADH/NAD^+^ ratio. Transcriptomics data shows that upon Fe^3+^ depletion, *Mtb* down-regulates the operon encoding ATP synthase and the type I NADH dehydrogenase (**Dataset S1** and Kurthkoti et al. 2017; Serafini et al. 2013). We examined the levels of ATP and the NADH/NAD^+^ ratio in H37Rv and Erdman strain. Surprisingly, ATP levels (**Figure 1G and H**, **Figure 1–figure supplement 1 A and B**) did not decrease upon Fe^3+^ depletion and showed a trend very similar to the condition with sufficient Fe^3+^. In contrast, we did see a significant increase in the NADH/NAD^+^ ratio upon Fe^3+^ depletion compared to the presence of sufficient Fe^3+^ (**Figure 1I**, **Figure 1–figure supplement 1 C and D**), indicating an accumulation of NADH. The concordant results between the two wild type stains, supports the hypothesis that the loss of viability in Erdman is not due to cell death, and indicates that in *Mtb* Fe^3+^ starvation causes a metabolic and physiological stress that is distinct from that observed in hypoxia and nutrient starvation (Rao et al. 2008; Gengenbacher et al. 2010; Eoh & Rhee 2013).

### Slowdown of the Krebs cycle activity

Most genes encoding enzymes involved in the Krebs cycle exhibit a reduction in their expression levels over a two-week period in Fe^3+^-starved H37Rv (**Dataset 1** and Kurthkoti et al. 2017). This finding indicates that the activity of this pathway may be reduced in such a condition. The levels of Krebs cycle intermediates were examined following an eight-day exposure to Fe^3+^-replete conditions (50 μM FeCl_3_, termed HI henceforth) and Fe^3+^-limiting conditions (0 μM FeCl_3_ and 0 μM FeCl_3_+DFO, termed LI and DFO, respectively, henceforth). As the metabolomics analysis demonstrated highly comparable outcomes in the LI and DFO conditions, the subsequent sections will focus on the comparison between HI and DFO. Please refer to supplementary data (***Supplement file* section I**) for a detailed analysis in the LI condition.

In H37Rv, a consistent increase, exceeding a 100-fold change (FC), in the intracellular levels of pyruvate and α-ketoglutarate was observed in DFO compared to HI (**Figure 2A**). The accumulation of these two metabolites is in line with the downregulation of the *aceE*, *dlaT* and *lpdC* genes (**Dataset S1** and Kurthkoti et al. 2017), which encode subunits of the pyruvate dehydrogenase (PDH) and the α-ketoglutarate dehydrogenase (KDH) complexes (Tian et al. 2005; Shi & Ehrt 2006; Maksymiuk et al. 2015). This accumulation is consistent with the existence of two stalling points at which the activity of the Krebs cycle is impeded, one before the cycle (PDH) and one at the KDH level. It is likely that the blockage at the level of pyruvate results in a slowdown the flux through glycolysis. The downregulation of glycolytic genes (**Dataset 1** and Kurthkoti et al. 2017) lends support to this hypothesis. A smaller increase in the levels of (iso)citrate was observed, with a 2- to 4-FC in two independent experiments (**Figure 2A**). This increase may be attributed to two potential mechanisms: i) citrate accumulation resulting from the reduced activity of the iron-dependent aconitase (Acn), which is also known to be down-regulated (Kurthkoti et al. 2017; Wong et al. 1999; Serafini et al. 2013); ii) isocitrate accumulation due to blockage of the Krebs cycle at the KDH level. The major fold increase (∼100-fold) in αKG compared to (iso)citrate is consistent with the irreversibility of the isocitrate dehydrogenase reaction, which drives carbon flux from isocitrate toward αKG but not in the reverse direction. As a result, isocitrate cannot accumulate to the same extent. Further, the intracellular threshold of these two metabolites may differ leading the cells to divert (iso)citrate into other metabolic pathways, limiting its accumulation.

**Figure 2.**
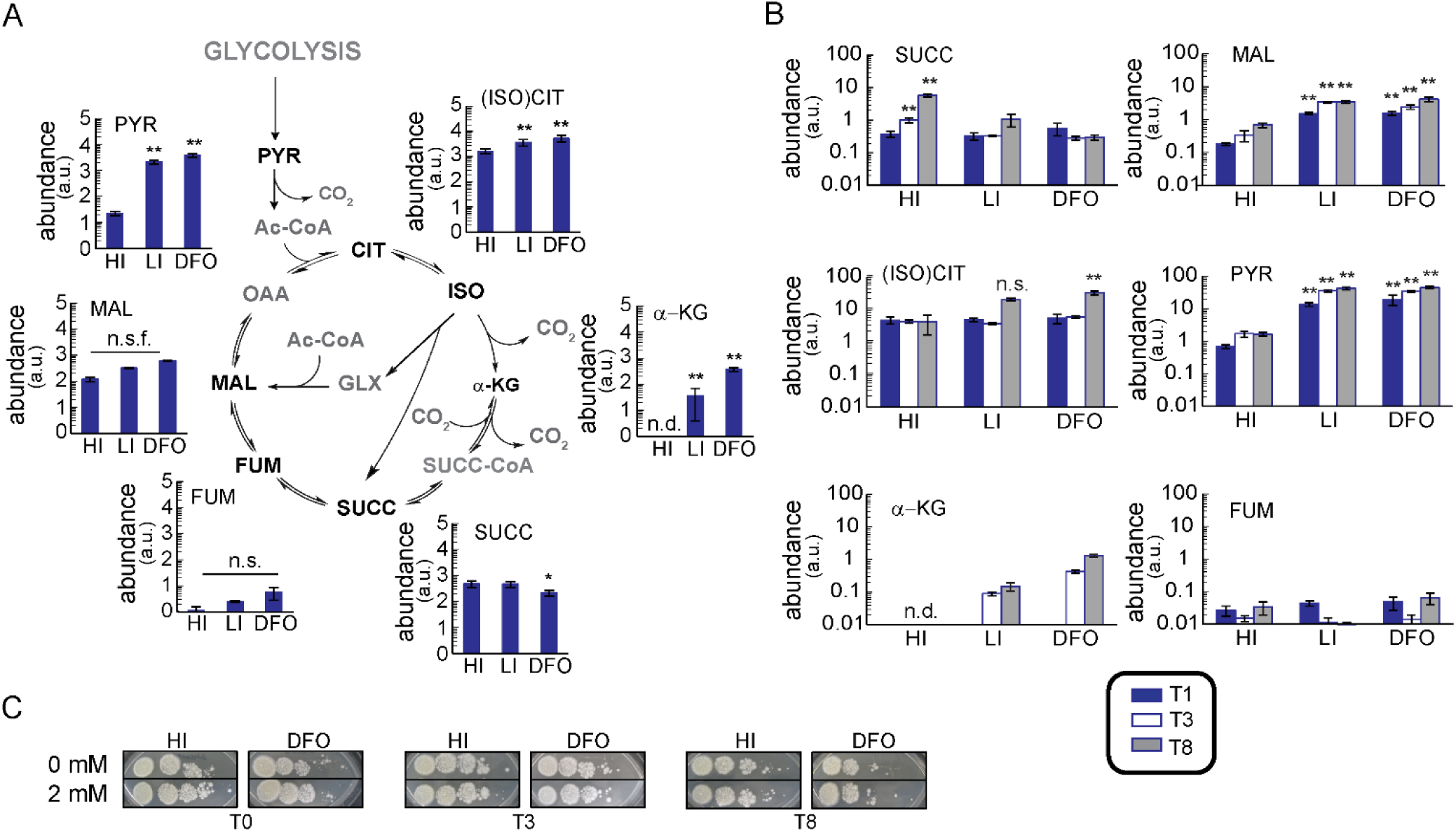
Intracellular and extracellular levels of metabolites in H37Rv. Cells were exposed to 50 μM FeCl_3_ (HI: High Iron), 0 μM FeCl_3_ (LI: Low Iron,) or 0 μM FeCl_3_+ DFO (DFO). The analysed metabolites are shown in black in the schematic pathways. A) Intracellular polar metabolites levels at 8 days; the y-axis is shown on Log_10_ scale tick labels (0–5) represent the exponents of 10 (10⁰–10⁵); values are reported in arbitrary units. B) Extracellular polar metabolites levels at 1, 3 and 8 days. The plots show the normalised levels of metabolites. The data represent the average and the standard deviation of four biological replicates from an independent experiment, representative of two independent experiments. The y-axis is shown on is in Log_10_ scale, values are reported in arbitrary units. C) Viability of H37Rv from one independent experiment. Cells were grown in liquid medium in HI and DFO conditions and in the presence of 2 mM of succinate. Aliquots of cells were collected after 0, 3 and 8 days and diluted to same final OD_600_; 5 μL of 10-fold serial dilutions were plated on 7H10. Growth was recorded after 19-25 days. The p-values were calculated against the HI condition and independently for the two experiments, the highest p-value was reported. n.s.f.= non-significant fold change, the observed trend change was different between independent experiments; n.s.= non-significant, p value >0.05; n.d.= non-detected; * =p value <0.05; ** =p value <0.01. Ac-CoA: acetyl-CoA; CIT: citrate; FUM: fumarate; α-KG: α-ketoglutarate; ISO: isocitrate; (ISO)CIT: isocitrate and citrate; MAL: malate; OAA: oxaloacetate; PYR: pyruvate; SUCC: succinate; SUCC-CoA: succinyl-CoA.

The succinate pool exhibited a statistically significant decrease in DFO, with intensity varying from 0.3- to 0.8-FC in two independent experiments (**Figure 2A**). No relevant alterations were observed in the levels of fumarate and malate. These results seem to contrast with the observation that genes encoding enzymes involved in the synthesis of fumarate and malate (fumarate reductase/Frd, fumarase/Fum and succinate dehydrogenase/Sdh) are iron-dependent and downregulated under Fe^3+^ starvation (**Dataset S1** and Kurthkoti et al. 2017) and therefore their activity should be lower in such conditions. Surprisingly, the genes encoding the α-ketoglutarate ferredoxin-oxidoreductase KorAB complex (Baughn et al. 2009) are induced (Kurthkoti et al. 2017) suggesting that despite its iron-dependent nature, its activity is necessary to produce succinyl-CoA and then succinate from α-ketoglutarate.

As observed in H37Rv, in the Erdman strain (**Figure 2–figure supplement 1A**) there is a substantial intracellular accumulation of pyruvate and α-ketoglutarate (>10 FC) and a more modest accumulation of (iso)citrate (1.2-2.8 FC, in four independent experiments) in DFO. No major changes were observed in the levels of malate, fumarate and succinate.

In conclusion, when experiencing Fe^3+^ starvation, *Mtb* slows down the activity of the Krebs cycle, and pyruvate conversion to acetyl-CoA and α-ketoglutarate conversion into succinyl-CoA may represent two distinct checkpoints in this process.

### Substantial secretion of metabolic intermediates

The secretion of succinate has been linked to the maintenance of membrane potential in hypoxic growth-arrested *Mtb* (Eoh & Khee 2013). It has been demonstrated that in such condition the extracellular succinate levels increase and that the inhibition of its secretion (by supplementing the medium with succinate) causes alteration of the proton motive force and cell death (Eoh & Khee 2013). We hypothesised that succinate could have a similar role in DFO-treated cells (Serafini 2021). Levels of extracellular succinate were therefore monitored over a week period (1, 3 and 8 days). Unexpectedly, no change in succinate levels was observed in DFO, rather, a significant increase was noted in HI (**Figure 2B and Figure 2–figure supplement 1B**). Furthermore, the succinate levels were lower in DFO compared to the HI condition. Since *Mtb* cells remain viable in DFO (**Figure 1C and D**), it can be concluded that the secretion of succinate is not a crucial factor in maintaining cell survival and membrane potential in such condition. This conclusion was confirmed by the observation that DFO-treated cells exposed to 2 mM succinate in the order to inhibit its secretion (Eoh & Khee 2013), remained viable (**Figure 2C and Figure 2-figure supplement 1C**), in contrast to the results observed during hypoxia.

Surprisingly, higher levels of several Krebs cycle intermediates were found in the culture filtrate of DFO compared to HI cultures, with minor discrepancies between the H37Rv (**Figure 2B)** and Erdman (**Figure 2–figure supplement 1B**) strain. α-ketoglutarate, (iso)citrate, malate and pyruvate are secreted in substantial quantities and their levels increase over time, with the exception of fumarate and succinate. The levels of pyruvate and α-ketoglutarate increase by more than 10-fold, while those of (iso)citrate rise by 2- to 10-fold under Fe^3+^ starvation. The high levels of extracellular pyruvate, α-ketoglutarate and (iso)citrate are consistent with the elevated intracellular levels. It is noteworthy that malate levels exhibit a substantial increase, approximately 10-fold, in DFO compared to HI conditions despite similar intracellular levels under the two conditions. This increase occurs after one day of DFO exposure in H37Rv (**Figure 2B**) and at a later stage in the Erdman strain (**Figure 2-figure supplement 1B**). The analysis of ^13^C incorporation in extracellular metabolites is consistent with the altered flow of carbon through the Krebs cycle (**Figure 2-figure supplement 2A-H** and **Supplement file S*ection II***).

The decrease in CFU/mL in Erdman (**Figure 1D**) raised the possibility of cell lysis, that could explain the increase of extracellular malate despite of unchanged intracellular levels in DFO compared to HI conditions. If increased lysis had occurred, we would expect an overall increase in extracellular metabolites. However, extracellular fumarate and succinate levels did not increase in DFO both in Erdman strain and H37Rv (**Figure 2B and Figure 2-figure supplement 1B**). We also examined the levels of extracellular glutamate, an amino acid which abundance increase over a week in HI. As for fumarate, extracellular glutamate levels remained unchanged or decreased in DFO (**Figure 2-figure supplement 2 I and L**) in Erdman and H37Rv. Therefore, we conclude that the observed increase of abundance of specific metabolites is not due to cell lysis.

As the intracellular malate pool is constant under HI and DFO conditions, this increase in malate secretion suggested that this metabolite may be utilised for the maintenance of membrane potential. The viability of DFO-treated cells in the presence of malate was evaluated (**Figure 2-figure supplements 1C and 3A**) but no effect was observed on survival. The substantial secretion of several metabolites prompted the hypothesis that they may collectively contribute to the maintenance of the membrane potential. The experiment was then repeated adding pyruvate, α-ketoglutarate, succinate, malate and fumarate individually (**Figure 2-figure supplements 1C and 3A**) and in combination, yet again we saw no loss of viability. These results suggest that the secretion of these metabolites under Fe^3+^ starvation is unrelated to the maintenance of the membrane potential.

### Partitioning of carbon flux into oxidative and reductive branches of Krebs cycle

The transcriptomic data from Fe^3+^-starved H37Rv (Kurthkoti et al. 2017; Serafini et al. 2013), shows upregulation of isocitrate lyase 1 (*icl1/Rv04671*) (Honer Zu Bentrup et al. 1999) and phosphoenolpyruvate carboxykinase A (*pckA*/*Rv0211*) (Marrero et al. 2010; Machova et al. 2015) genes (**Dataset S1**). To verify the effective operation of these metabolic routes, cells were fed with ^13^C_3_-glycerol and the isotopic labelling profiles of multiple metabolites reporting on CCM were analysed (**Figures 3 and 4**).

**Figure 3.**
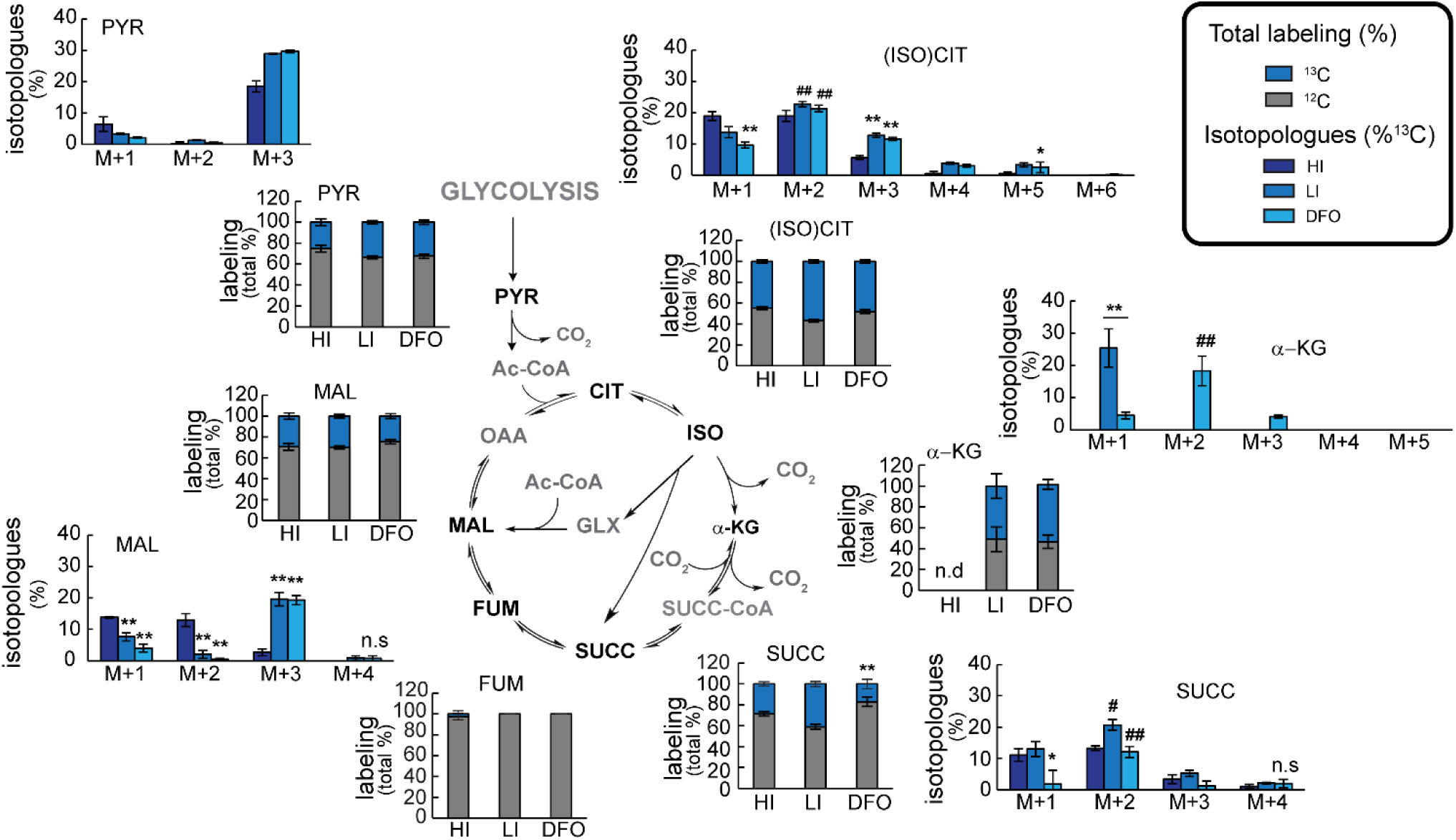
Percentage of labelled (^13^C) and unlabelled (^12^C) metabolites in H37Rv. Metabolites were extracted from cells fed with ^13^C_3_-glycerol for 8 days in 50 μM FeCl_3_ (HI: High Iron), 0 μM FeCl_3_ (LI: Low Iron) or 0 μM FeCl_3_+ DFO (DFO). The analysed metabolites are shown in black in the schematic pathways. For each metabolite, but fumarate, two plots are shown. The stacked column plot shows the total percentage of labelled and unlabelled molecules per each metabolite pool; the clustered column plot shows the abundance in percentage of each isotopologue. The data represent the average and the standard deviation of four biological replicates from an independent experiment, representative of two independent experiments. The p-values were calculated independently for the two experiments and the highest p-value was reported. DFO/LI vs HI condition: * = p value<0.05; ** = p value <0.01; n.s.= non-significant. n.d.= non-detected. M2 vs M1: # = p value<0.05; ## = p value<0.01. Ac-CoA: acetyl-CoA; CIT: citrate; FUM: fumarate; α-KG: α-ketoglutarate; ISO: isocitrate; (ISO)CIT: isocitrate and citrate; MAL: malate; OAA: oxaloacetate; PYR: pyruvate; SUCC: succinate; SUCC-CoA: succinyl-CoA.

**Figure 4.**
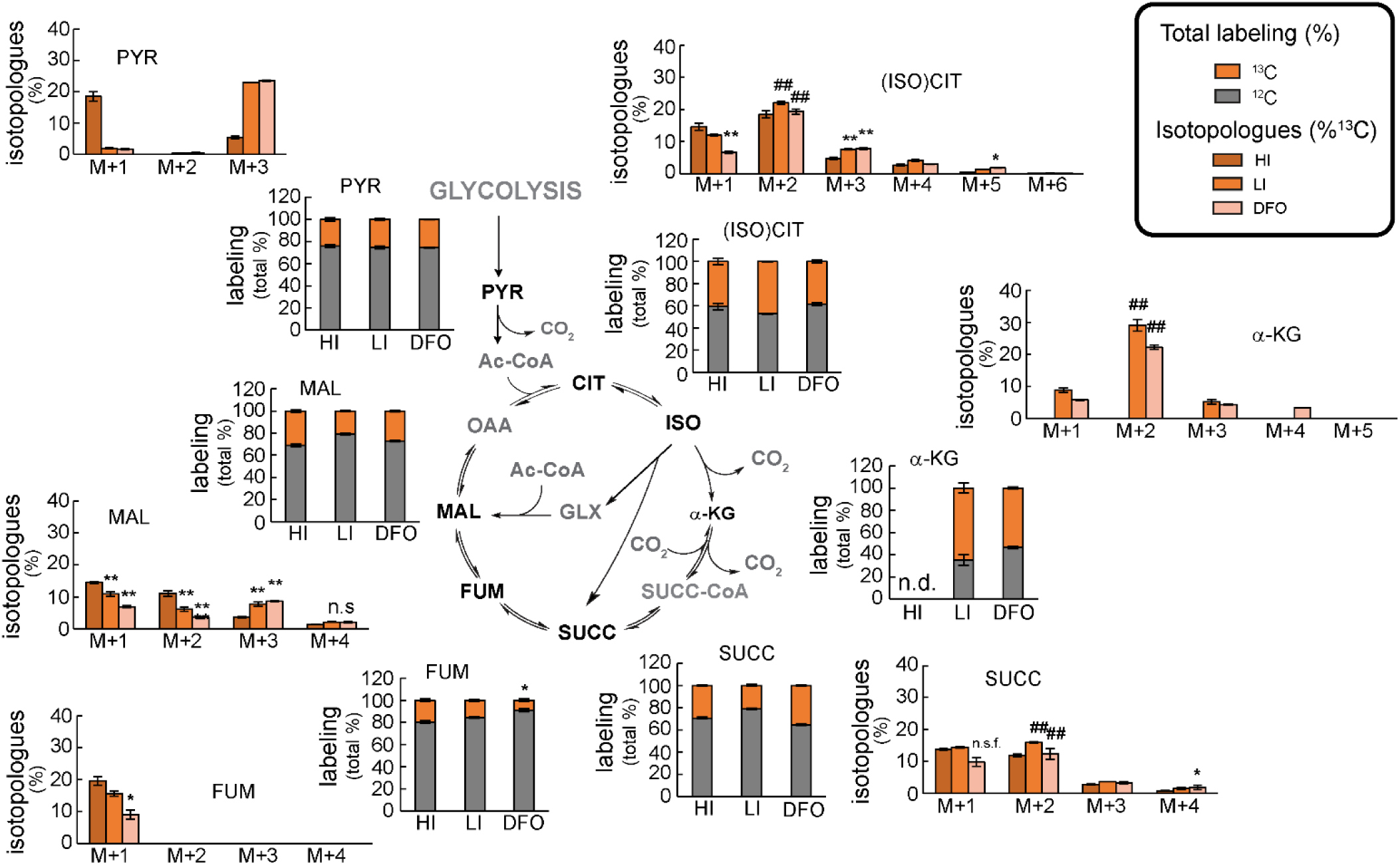
Percentage of labelled (^13^C) and unlabelled (^12^C) metabolites in the Erdman strain. Metabolites were extracted from cells fed with ^13^C_3_-glycerol for 8 days in 50 μM FeCl_3_ (HI: High Iron), 0 μM FeCl_3_ (LI: Low Iron) or 0 μM FeCl_3_+ DFO (DFO). The analysed metabolites are shown in black in the schematic pathways. For each metabolite two plots are shown. The stacked column plots show the total percentage of labelled and unlabelled molecules per each metabolite pool; the clustered column plots show the abundance in percentage of each isotopologue. The data represent the average and the standard deviation of four biological replicates from an independent experiment, representative of four independent experiments (only two for LI condition). The p-values were calculated independently between experiments and the highest value is reported. <u>DFO/LI vs HI</u>: ** = p value <0.01; n.s.= non-significant; n.s.f.= non-significant fold change, the observed trend change was different between independent experiments. n.d.= non-detected. <u>M2 vs M1</u>: ## = p value <0.01. Ac-CoA: acetyl-CoA; CIT: citrate; FUM: fumarate; α-KG: α-ketoglutarate; ISO: isocitrate; (ISO)CIT: isocitrate and citrate; MAL: malate; OAA: oxaloacetate; PYR: pyruvate; SUCC: succinate; SUCC-CoA: succinyl-CoA.

#### Malate only partially derives from succinate oxidation

DFO-treated H37Rv shows a 50% reduction in labelled succinate compared to HI condition (**Figure 3**). This decrease is not observed in isocitrate and α-ketoglutarate, precursors of succinate. These discrepancies suggest that a slower rate conversion of α-ketoglutarate into succinyl-CoA and subsequently into succinate, and of isocitrate to succinate may occur in DFO. The down-regulation of KDH complex genes (**Dataset S1** and Kurthkoti et al. 2017) supports the hypothesis related to α-ketoglutarate, however the up-regulation of *icl1* and *korAB* (see above) contradicts both hypotheses. The percentage of labelling observed in the malate pool (**Figure 3**) remains constant between the DFO and HI conditions. Notably, the percentage of malate labelling is significantly higher than that of succinate in DFO, indicating that some of the malate may not originate from succinate via fumarate. Unfortunately, the fumarate levels (**Figure 3**) were insufficient to permit the detection of labelling for this metabolite and verify if it is similar to malate rather than succinate. However, the comparison of the isotopologue distributions of malate and succinate confirms that some of the malate is not a product of succinate oxidation (**Figure 3**). Indeed, whereas in HI conditions the malate and succinate labelled pools show similar levels of M+1 and M+2 isotopologues (omitted isotopologue word after ‘M+*n*’ henceforth) and lower levels of M+3, in DFO M+2 succinate is more abundant compared to the M+1 and M+3, and M+3 malate is the most abundant (70 - 90% of total labelling) over M+1 and M+2 (**Figure 3**).

The DFO-treated Erdman strain does not show a marked decrease in the total labelling of succinate pool (**Figure 4**), and the isotopologue distribution shows that M+1 and M+2 are present at similar levels in the DFO and HI conditions. Similarly to H37Rv, in Erdman strain the malate labelled pool size was similar in DFO and HI conditions, and contains more M+3 in DFO compared to HI (40 – 50% of total labelling), with a clear decrease in M+1 an M+2 (**Figure 4**). In the Erdman strain, higher levels of fumarate were detected and ^13^C incorporation could be evaluated. Surprisingly, the labelled fumarate pool contained only M+1 in all conditions (**Figure 4**), impeding a more precise tracking of its origin. This specific labelling profile of fumarate is likely linked to the specific medium used in the current work (**Supplement file section III**).

Altogether, these results strongly suggest that up to 50% of the malate pool is not derived from succinate oxidation under the DFO condition in *Mtb*.

#### Malate mainly derives from oxaloacetate reduction

The upregulation of *icl1* and *pckA* genes under Fe^3+^ condition (Kurthkoti et al. 2017; Serafini et al. 2013) suggests that malate may be produced via isocitrate lyase (ICL) and phosphoenolpyruvate carboxykinase (PCK) activities, despite the conflicting differences in the total labelling (**Figure 3**) between isocitrate and succinate (see above). The increase of these enzymatic activities was confirmed in the cell-free extracts (**Figure 5A and B**, **Figure 5-figure supplement 1**).

**Figure 5.**
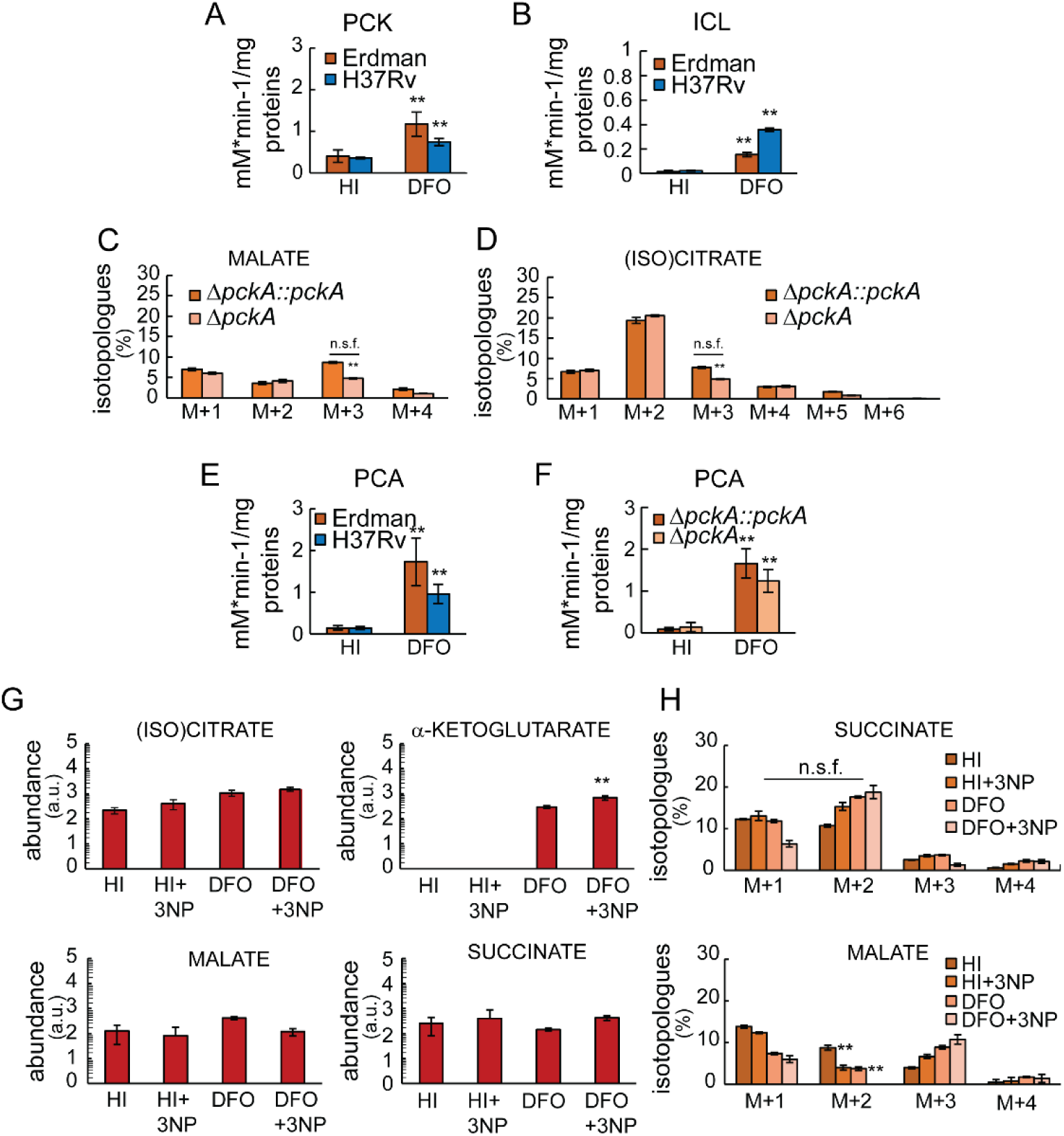
Analysis of iron-independent metabolic routes. Cells were exposed to 50 μM FeCl_3_ (HI), or 0 μM FeCl_3_+ DFO (DFO). A) Enzymatic activity of phosphoenolpyruvate carboxykinase (PCK), reaction from phosphoenolpyruvate to oxaloacetate; B) Enzymatic activity of isocitrate lyase (ICL); the plots show the activity in (mM of NADH × min^-1^)/mg of total protein detected in cell-free extracts after 3 days of exposure to DFO or HI condition. Data show average and standard deviation from two (H37Rv) or three (Erdman strain) independent experiments and two technical replicates each (A) or from one independent experiment and three technical replicates (B). C, D) Isotopologue distribution of intracellular malate (C) and isocitrate (D) in Erdman-derived *ΔpckA::pckA* and *ΔpckA* strains after 8 days of exposure to DFO or HI conditions and fed with ^13^C_3_-glycerol. The plots show the abundance in percentage of each isotopologue. The histograms represent average and standard deviation from four biological replicates from one experiment, representative of three independent experiments. E, F) Enzymatic activity of pyruvate carboxylase (PCA), reaction from pyruvate to oxaloacetate; the plots show the activity in (mM of NADH × min^-1^)/mg of total protein detected in cell-free extracts after 3 days of exposure to DFO or HI condition. Data show average and standard deviation from three independent experiments and two technical replicates each. G, H) Erdman strain cells were exposed to HI or DFO condition for 8 days with or without 200 μM of inhibitor 3-nitropropionate (3NP) and fed with ^13^C_3_-glycerol. (G) The plots show the normalised levels of metabolites; the y-axis is shown on Log_10_ scale tick labels (0–5) represent the exponents of 10 (10⁰–10⁵); values are reported in arbitrary units. (H) The plots show the isotopologue distribution in percent abundance. The data represent the average and the standard deviation of four biological replicates from an independent experiment, representative of two independent experiments. The p-values were calculated independently between experiments and the highest value is reported. The p-values were calculated as follows. <u>DFO vs HI</u> for A, B, E, F; <u>HI+3NP vs HI</u> and <u>DFO+3NP vs DFO</u> for H; <u>mutant vs complemented</u> for C, D. n.s.f. = non-significant fold change, the observed trend change was different between independent experiments. ** = p value <0.01.

**Figures 3 and 4-figure supplements 1 and 2** show 31 possible metabolic scenarios for the production of core of CCM intermediates when the glyoxylate cycle and/or the PCK anaplerotic route are active. These scenarios combine labelled and unlabelled pyruvate, phosphoenolpyruvate (PEP), acetyl-CoA and carbon dioxide molecules. Although they do not consider the re-circulation of metabolites in the global metabolic circuit, these series of reactions offer a useful perspective on identifying operating pathways. We compared these theoretical possibilities with our labelling data (isotopologue distribution) on succinate, malate, and (iso)citrate (**Figures 3 and 4**). Oxaloacetate was not included in this analysis due to technical limitations (**Supplement file Section IV**). Of note, the isotopologue distribution of succinate is identical whether derived from isocitrate or from α-ketoglutarate.

**Figure 3 and 4-figure supplement 1A-C** show scenarios derived from pyruvate dehydrogenase (PDH) and ICL activities. M+2 (iso)citrate, malate and succinate are the only isotopologues produced. This matches the experimental results for isocitrate and succinate, for which M+2 is one of the most abundant isotopologues (**Figure 3 and 4**). Of note, the medium used in these experiments contains asparagine, which represents an exogenous unlabelled source of oxaloacetate (**Supplement file section IV**). This indicates that the **Figure 3 and 4-figure supplement 1A-C** scenarios are favoured and explain the dominant abundance of M+2 (iso)citrate and succinate, compared to other isotopologues. However, these combinations do not produce the most abundant malate isotopologue, M+3, detected in our measurements.

**Figure 3 and 4-figure supplement 1D-S** show PCK and ICL activities. In these scenarios the M+3 malate isotopologue is present and derives from the glyoxylate shunt. However, there is no congruence between these theoretical scenarios and our labelling results (more details in **Supplement file section V**). This raises doubts about an active role of the glyoxylate shunt in the synthesis of malate.

**Figure 3 and 4-figure supplement 2A-H** illustrate alternative metabolic scenarios without the involvement of the glyoxylate shunt that include PCK activity and the oxidative branch of the Krebs cycle stopping at malate. Among these scenarios, the most represented isotopologues are M+3 for (iso)citrate (**Figure 3 and 4-figure supplement 2B and G**), M+2 for malate and for succinate (**Figure 3 and 4-figure supplement 2B, C, F and G**). These scenarios offer a rationale for the observed increase in M+3 (iso)citrate in our experiments in DFO (Figure 3 **and 4**), and imply that M+2 malate and succinate may originate from the oxidative branch of the Krebs cycle, fed via oxaloacetate produced by PCK (**Figure 3 and 4-figure supplement 2B, C, F and G**). These alternative scenarios align with those showed in **Figure 3 and 4-figure supplement 1A-C** for the synthesis of M+2 malate and succinate, excluding the glyoxylate shunt activity, but do not reveal the origin of M+3 malate.

We then hypothesised that PEP-derived oxaloacetate could proceed to the reductive branch of Krebs cycle (**Figure 3 and 4-figure supplement 2I-L**). The malate isotopologues produced in the **Figure 3 and 4-figure supplement 2I, K and L** scenarios are all detected in our experiments. The dominance of M+3 over M+1 (Figure 3 **and 4**) indicates that the scenario illustrated in **Figure 3 and 4-figure supplement 2I** is the most likely. The minor abundance of M+4 malate (Figure 3 **and 4)** is compatible with the unlikely possibility that an enzyme meets both substrates labelled (**Figure 3 and 4-figure supplement 2K**). A similar consideration is valid for the generation of M+4/M+5/M+6 (iso)citrate and M+4 succinate (**Figure 3 and 4-figure supplement 2A, E and F,** Figures 3 and **4**). This new analysis, which omits the glyoxylate shunt, is consistent with our experimental results and indicates that M+2 and M+4 malate derive from the oxidative branch of Krebs cycle, while M+1 and M+3 malate from the reductive branch. Further, these results indicate that, *pckA* has an important role in the CCM under Fe^3+^ starvation. Surprisingly, an Erdman-derived *pckA* mutant, which does not show loss of viability and reducing power under exposure to Fe^3+^ starvation (Figure 1F and I), does not show a significant decrease in M+3 malate and M+3 (iso)citrate in DFO (Figure 5C and D). In fact, there is no consistent between experiments (0-50% of decrease). Oxaloacetate can be produced directly from pyruvate by a pyruvate carboxylase (PCA) (Basu et al. 2018; Mukhopadhyay & Purwantini 2000), generating the same scenarios depicted in **Figure 3 and 4-figure supplement 1 and 2**, or malate can be directly produced from pyruvate by malic enzyme (MEZ) (**Figure 3 and 4-figure supplement 2M-P**). Measurement of the activity of these two enzymes in cell free extracts revealed the presence of PCA activity in the DFO condition (Figure 5 **E and F,** Figure 5**-figure supplement 1**), but not of MEZ activity (Figure 5**-figure supplement 1**). The absence of *pckA* does not result in an increase of PCA activity (Figure 5F), suggesting that the two activities work in parallel, but serve distinct and non-redundant functions. These results strongly suggest that *Mtb* utilises anaplerotic reactions mediated by PCK and PCA to supply Krebs cycle intermediates under Fe^3+^ starvation (**Figure 3 and 4-figure supplement 2I**).

### The glyoxylate shunt does not supply malate and succinate under iron starvation

To provide independent, genetic evidence supporting or ruling out the role of the glyoxylate shunt under Fe^3+^ starved conditions in H37Rv, the levels of succinate, malate and isocitrate were analysed in an isocitrate lyase-deleted strain. H37Rv has two ICL enzymes, encoded by the *icl*1 and *aceAab* genes, however the second enzyme has a slower activity (Honer Zu Bentrup et al. 1999) and studies show that the inactivation of just *icl1* is relevant to alter *Mtb* physiology (Serafini et al. 2019; Gengenbacher et al. 2010). We used an H37Rv-*Δicl1* and its complemented strain (Serafini et al. 2019), which is not essential for survival in DFO condition (Figure 1E). As the *Δicl1* strain is not derived from the same H37Rv strain used in this study, metabolite abundance and isotopologue distribution in HI, LI and DFO conditions were verified and confirmed in the complemented strain (*Δicl1::icl1*). Except for the lack of increase in intra- and extracellular (iso)citrate levels, all other trends were matched (**Figure 2 and 3-figure supplement 1**). If the glyoxylate shunt is a key supplier of malate and succinate, the absence of a functional ICL should result in a reduction in their abundance or labelling, an increase in isocitrate levels and a decrease in the M+2 in succinate. To our surprise, there are no discernible differences between the *Δicl1* and the complemented strains, in the intracellular levels of these metabolites (**Figure 2 and 3-figure supplement 2A**) and in the isotopologue distribution of succinate and malate (**Figure 2 and 3-figure supplement 2B and C**) in DFO condition. We reasoned that in *Δicl1* metabolite secretion could be diminished to maintain the intracellular abundance of malate and succinate or increased in the case of isocitrate. Again, there are no significant differences in the extracellular levels of these metabolites between the *Δicl1* and the complemented strains (**Figure 2 and 3-figure supplement 2E**).

Next, we verified the operation of glyoxylate shunt in the Erdman strain by analysing the levels and labelling profiles of intracellular metabolites in DFO-treated cells exposed to a potent inhibitor of mycobacterial ICL activity, 3-nitropropionate (3NP) (Honer Zu Bentrup et al. 1999; Moynihan & Murkin 2014), which does not affect survival under such condition (Figure 2**-figure supplement 3B**). The presence of 3NP did not increase the abundance of (iso)citrate, but rather that of α-ketoglutarate, and no relevant changes were detected in malate and succinate levels (Figure 5G). The isotopologue distribution analysis revealed no change in the abundance of the M+2 succinate in the DFO condition, but rather a diminution of the M+2 malate (Figure 5H). 3NP is also an inhibitor of Sdh (Alston et al. 1977), and the observed result appears to report on that, instead of on inhibition of ICL. The results demonstrate that the glyoxylate shunt is not playing an important role under Fe^3+^ starvation in *Mtb*. However, the presence of isocitrate lyase activity in lysates indicates that the enzyme is active and could generate a pool of succinate, yet not detectable under our experimental conditions, likely due to compensatory metabolic routes (see below). The labelling profile analysis (Figure 3 **and 4**) excludes malate as a derivative of glyoxylate. We therefore investigated an alternative metabolic fate of this molecule. Glyoxylate can be converted to glycine via reductive amination catalysed by alanine dehydrogenase (Ald), whose transcript is upregulated under iron limitation (Griffin et al., 2012; Kurthkoti et al., 2017) leading us to hypothesise that this metabolic route might be active under iron starvation. Due to technical limitations, glyoxylate could not be directly detected, but we assessed glycine abundance and its isotopologue distribution (Figures 3 and 4-fig**ure supplement 3 A-D**), which should mirror that of glyoxylate if glycine is derived from it. While total glycine levels were unchanged between HI and DFO conditions, isotopologue analysis revealed a ∼3-fold increase in M+2 relative to M+1 under DFO in both H37Rv and Erdman strains. The comparison (Figures 3 and 4-fig**ure supplement 3E**) of these results with the most probable labelling scenarios of glyoxylate illustrated in Figures 3 and 4-fig**ure supplements 1 and 2**, suggests that glycine does not originate from glyoxylate. Additionally, no differences in labelling were observed between the *icl1* mutant and the complemented strain (Figures 3 and 4-fig**ure supplement 3 F and G**). Taken together, these results indicate that glyoxylate is not a precursor of glycine under iron starvation, further supporting the non-operational state of the glyoxylate shunt under these conditions.

The lack of evidence for an operational glyoxylate shunt under Fe^3+^ starvation led us to conclude that succinate must derive from α-ketoglutarate oxidation. The upregulation of the *korAB* genes (Kurthkoti et al. 2017) suggests that KorAB actively participates in succinyl-CoA synthesis. However, its iron-dependent nature together with the downregulation of succinyl-CoA synthetase genes (*sucCD*) suggests that an additional pathway, likely iron-independent, might be active under Fe^3+^ starvation conditions to maintain the succinate pool. This pathway might be the γ-aminobutyric acid (GABA) shunt, an active route of CCM in Mtb (Serafini et al. 2019), which converts α-ketoglutarate into succinate bypassing succinyl-CoA synthesis (Figure 5**-figure supplement 2A**). Genes encoding GABA shunt enzymes are not differentially expressed under Fe^3+^ starvation conditions (**Dataset S1** and Kurthkoti et al. 2017; Serafini et al 2013). In H37Rv, the levels and labelling of glutamate do not change to a major extent under HI and DFO conditions (Figure 5**-figure supplement 2B**). In contrast, we see a significant decrease in the GABA pool size in DFO (0.001 - 0.428 FC in four independent experiments), with a concomitant decrease in total labelling (reduction >50%) (Figure 5**-figure supplement 2C**). Changes in GABA abundance resemble that of succinate (Figure 5**-figure supplement 2D),** with a greater magnitude. This suggests that GABA might be used to produce succinate. We wondered if this reduction of the GABA pool could be due to its secretion, however we were not able to detect extracellular GABA (data not shown). The isotopologue distribution of the GABA and succinate pools (Figure 5**-figure supplement 2 C and D**) are comparable to those of α-ketoglutarate (Figure 3) with a notable reduction in the M+1 compared to the M+2. Similar results were obtained in the Erdman strain (Figure 5**-figure supplement 2 E, F and G**) in which the isotopologue distribution of succinate resembles that of GABA (Figure 5**-figure supplement 2 F and G**) and glutamate, yet, it differs from that of α-ketoglutarate (Figure 4, Figure 5**-figure supplement 2E**). In particular, in succinate, GABA, and glutamate the levels of M+1 and M+2 are similar in DFO (Figure 5**-figure supplement 2E, F and G**), whereas M+2 α-ketoglutarate consistently exceeds M+1 (Figure 4). This different isotopologue distribution in α-ketoglutarate suggests that the GABA shunt is significantly sustained by other metabolic circuits beyond α-ketoglutarate in Erdman.

The absence of a differential expression of GABA shunt genes under Fe^3+^ starvation and the similarity between the labelling profiles of succinate and GABA are consistent with the hypothesis that the GABA shunt may contribute to maintaining the succinate pool under Fe^3+^ starvation conditions in *Mtb*, although direct genetic evidence is still required to substantiate this conclusion."

## Discussion

This study was conducted on two *Mtb* strains that exhibit similar behavior, with minor differences likely reflecting variations in the intensity of Krebs cycle activity (**SI Appendix, Section II**). The results of this investigation are summarised in Figure 7. The consistent accumulation of pyruvate and α-ketoglutarate highlights two key check points controlling carbon flux through the CCM. Because DlaT and LpdC are shared components of both the pyruvate and the α-ketoglutarate dehydrogenase complexes (Tian et al. 2005; Shi &Ehrt 2006; Maksymiuk et al.2015), they likely play a critical role in this buildup. As central metabolites linking energy and carbon/nitrogen metabolism, pyruvate and α-ketoglutarate accumulation indicate that Fe^3+^ starvation slows down the overall cellular metabolism. Under these conditions, *Mtb* cells survive and maintain a physiologic energy balance (Figure 1G and H **and** Figure 1**–supplement figure 1A and B**) but display an altered redox state marked by increased NADH levels (Figure 1**–supplement figure 1C and D)**. It has been observed that under Fe^3+^ starvation *Mtb* downregulates genes encoding the proton pumping enzymatic complexes Type I NADH dehydrogenase (*nuoABCDEFGHIJKLMN*) and cytochrome oxidase bc_1_-aa_3_ (*qcrCAB*/*ctaBCED*), whereas it upregulates the gene for non-proton pumping Type II NADH dehydrogenase (*ndh*) and the gene for assembly of the less efficient and non-proton pumping cytochrome oxidase *bd* (*cydABDC*) (Kurthkoti et al. 2017; Serafini et al 2013; Cruz-Ramoset al. 2004). The iron independent nature of Ndh and its lack of proton pumping activity may explain the opposite regulation of type I and II NADH dehydrogenases under Fe^3+^ starvation. In contrast, the reason behind the divergent regulation of the two cytochrome oxidases remains not clear. Together with the increased NADH/NAD ratio, these changes in the electron transport chain (ETC) suggest that the alternative ETC is less efficient at NADH reoxidation. Notably, the replacement of proton-pumping components with the non-proton pumping alternatives, along with the reported downregulation (Kurthkoti et al. 2017) of the ATP synthase operon (*atpBEFHAGDC*), contrasts with the stable ATP levels observed in Fe^3+^-starved *Mtb* cell.

Intriguingly, the differential expression of *atpBEFHAGDC, ndh,* and *nuoABCDEFGHIJKLMN* genes during Fe^3+^ starvation resembles that seen during the efficient *Mtb* growth in L-lactate as a sole carbon source (Serafini et al. 2019) suggesting that *Mtb* assembles an alternative ETC tailored to distinct physiological demands. Even more intriguing is the very recent work on *M. smegmatis*, which shows that a minimal ETC composed of cytochrome oxidase *b,* Ndh and F_1_F_0_-ATPase can generate ATP or *pmf* using atmospheric hydrogen (Soom et al. 2025). This raises the possibility that additional, yet unidentified mechanisms support ATP synthesis under the conditions used in our study.

The downregulation of transcripts for many CCM enzymes in Fe^3+^ starvation (Kurthkoti et al. 2017; Serafini et al 2013) is presumably a consequence of the slowed carbon flux, and the induction (Kurthkoti et al. 2017; Serafini et al 2013) of a few of them (*korAB*, *pckA* and *icl1*) is likely a strategy to sustain slowed carbon flow. The induction of the iron-dependent KorAB route represents a mechanism to bypass or parallel the KDH complex and GABA shunt, in order to sustain succinate synthesis and dispose of the accumulated α-ketoglutarate. This strategy combines multiple routes to maintain succinate synthesis under Fe^3+^-starved conditions. A combination of multiple routes is also deployed to sustain carbon flux from PEP/pyruvate to the Krebs cycle. The anaplerotic reaction of PCK converts PEP into oxaloacetate, and it is flanked by PCA, which converts pyruvate into oxaloacetate. These two activities, together with PDH, represent three routes to control the level of pyruvate and sustain the carbon flux. Detailed ^13^C tracking through Krebs cycle intermediates revealed an unexpected split of carbon flux from PEP- and pyruvate-derived oxaloacetate to both the oxidative and reductive branches of the Krebs cycle. Oxaloacetate is partitioned between oxidation to citrate and reduction to malate, with the latter route being the most favorable (Figures 3 **and 4**, Figure 3 **and 4-figure supplements 1 and 2**). The two fluxes converge to produce malate, the extracellular level of which increases (Figure 6). The production of oxaloacetate via PCK and PCA under Fe^3+^ starvation occurs despite the presence of an exogenous source of oxaloacetate (the asparagine in the medium, **Supplement file section IV**), and clearly demonstrates the importance of these anaplerotic reactions under Fe^3+^ starvation beyond oxaloacetate synthesis. These activities likely control pyruvate levels but also may contribute to assimilating the excess of carbon dioxide stoichiometrically produced with α-ketoglutarate by isocitrate dehydrogenase (Icd, Figure 6). The *pckA* gene is also induced in Fe^3+^ starvation when glutamate and ammonium, but not asparagine, are present as nitrogen sources in the medium (Serafini et al. 2013) suggesting that PCK and likely PCA are the main producers of oxaloacetate under these conditions.

**Figure 6.**
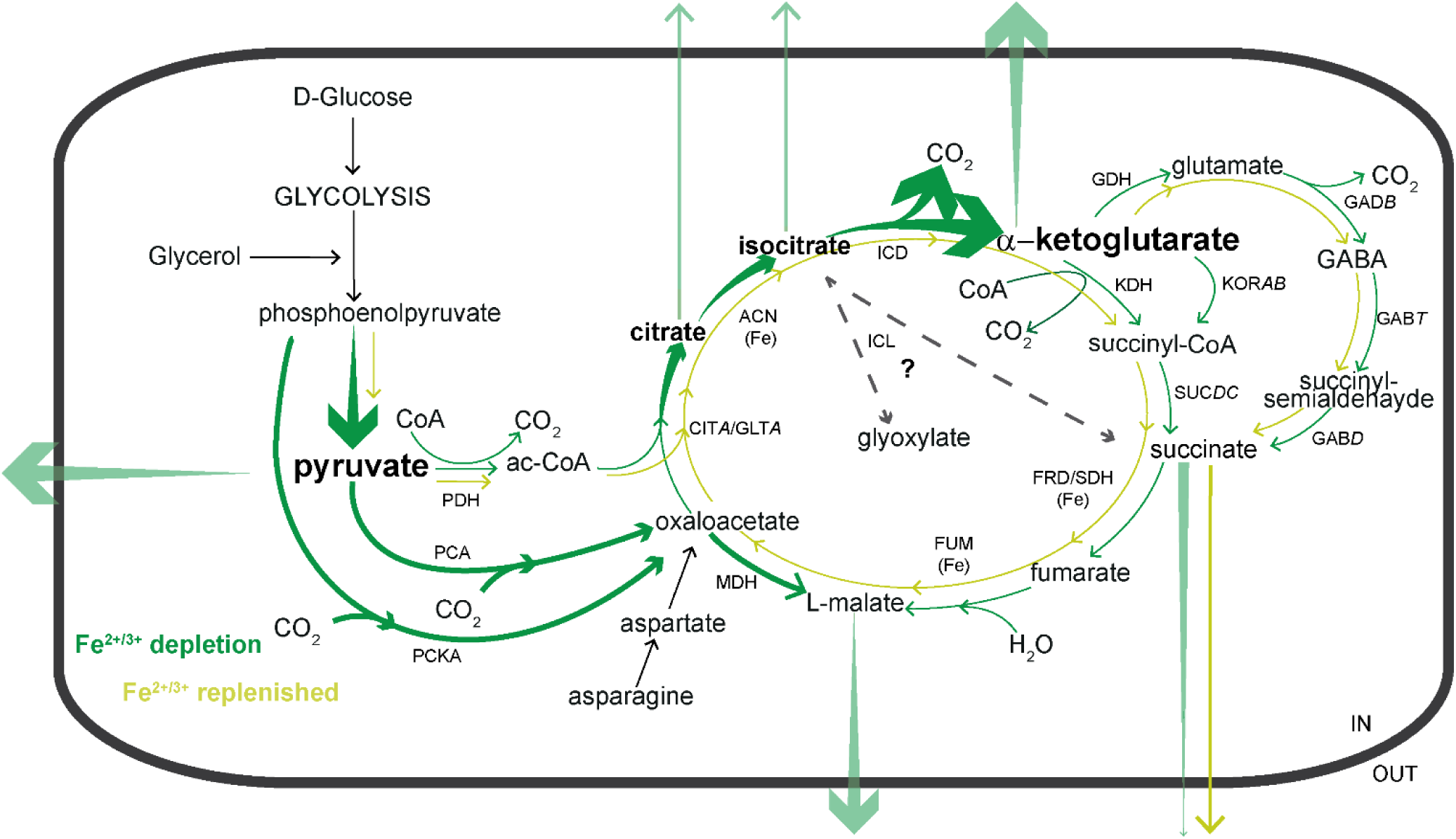
Re-modelling of CCM under iron starvation in *Mtb*. The picture depicts a schematic representation of the CCM pathways active in *Mtb* exposed to Fe^3+^ deprivation in the presence of D-glucose and glycerol as carbon sources and asparagine as sole nitrogen source. The thickening arrows indicate the increase of levels of the metabolite; thicker arrows indicate a preferred route. Bold and larger font indicates accumulated metabolites. Under iron starvation, the pool of iron-dependent enzymes (denoted by Fe in parentheses in the figure) contains a reduced number of fully active molecules, which then slows the carbon flux through the Krebs cycle. The reduction in the transcript levels of iron-independent enzymes of the CCM pathways is likely a consequence of this. The disparity in efficiency between iron-dependent and iron-independent enzyme pools gives rise to the accumulation of (iso)citrate, pyruvate, and α-ketoglutarate. *Mtb* responds to these accumulations by expelling these metabolites from the cell and splitting the carbon flux from PEP/pyruvate (via PCK, PCA, and PDH activities) into both oxidative and reductive branches of the Krebs cycle. Both fluxes terminate in malate synthesis, which is then secreted. To maintain malate synthesis by succinate oxidation *Mtb* limits its secretion. Malate secretion relieves the slowdown of carbon flux through the oxidative branch of Krebs cycle. PCK and PCA anaplerotic reactions control pyruvate levels and recycle carbon dioxide stoichiometric to the α-ketoglutarate accumulation. GDH: glutamate dehydrogenase. GAD*B*: glutamate decarboxylase. GAB*T*: 4-aminobutyrate aminotransferase. GAB*D*: succinate-semialdehyde dehydrogenase. For the other enzyme acronyms see main text.

The significant quantities of extracellular malate under Fe^3+^ starvation, despite its intracellular levels being similar to those in Fe^3+^ replete condition, suggests that secretion of this metabolite has a functional role. Also, the reduced secretion of succinate under Fe^3+^ starvation seems to indicate that succinate (Figure 2 **and** Figure 2**-supplement figure 1**) is required to produce malate. The secretion of malate may promote carbon flux through the anaplerotic node to relieve the accumulation of metabolic intermediates from the oxidative branch of the Krebs cycle. NADH re-oxidation mediated by malate dehydrogenase, together with Ndh, may help to maintain a redox state compatible with cell survival.

The upregulation of *icl1* (Kurthkoti et al. 2017; Serafini et al. 2013) and the resulting increase of ICL activity under Fe^3+^ starvation appear to be dispensable for the synthesis of malate and succinate (**Figure 2 and 3-supplement figure 2 and Figure 5**), which is surprising considering that the glyoxylate shunt has been demonstrated to be important for the survival of *Mtb* under various growth arrest-causing conditions (Eoh & Rhee 2013; Genegenbacher et al. 2010; Nandakumar et al. 2014). On the other hand, the key function of the glyoxylate shunt is to bypass the two Krebs cycle decarboxylation reactions, and thereby “save up” carbon. Under our experimental conditions, carbon is replete, and therefore the glyoxylate shunt is not needed. Isocitrate lyase might participate in a previously unrecognised metabolic pathway or other process. The induction of ICL under Fe^3+^ starvation is not unique to mycobacteria. An increase of its activity has been observed in *Pseudomonas aeruginosa* under Fe^3+^ starvation (Ha et al. 2018) and in *Pseudomonas fluorescens* under Al^3+^ stress, which mimics iron deficiency (Middaugh et al. 2005). In *P. aeruginosa*, the absence of ICL activity causes an increase in succinate dehydrogenase activity suggesting its involvement in succinate synthesis (Ha et al. 2018). It is noteworthy that *P. aeruginosa icl* mutant accumulates intracellular iron, suggesting a role beyond CCM. Unfortunately, metabolomics studies have not been performed in this mutant. Interestingly, it has been demonstrated that *P. fluorescens* exposed to Al^3+^ stress induces an acylating glyoxylate dehydrogenase (AGODH) that transforms glyoxylate to oxalyl-CoA and then oxalate, a molecule necessary to seize Al^3+^ ions (Singh et al. 2009). Coenzyme A is recycled to produce succinyl-CoA and then succinate, releasing one ATP (Singh et al. 2009). The finding that glyoxylate is used in a metabolic pathway functional for Al^3+^ stress, rather than for malate synthesis, supports the hypothesis that *Mtb* may use ICL for an alternative metabolic function under Fe^3+^ starvation.

Collectively, these findings suggest that, under iron depletion, *Mtb* relies on anaplerotic routes to divert carbon flux from glycolysis toward the reductive branch of the Krebs cycle. This results in a truncated cycle in which both carbon fluxes converge toward malate synthesis. The secretion of malate appears to be functionally relevant to sustaining these fluxes, ensuring the production of metabolic intermediates and maintaining redox balance through NADH reoxidation via malate dehydrogenase. Although the alternative ETC comprising Ndh and cytochrome oxidase *bd,* is less efficient in both ATP synthesis and NADH reoxidation, it may represent an adaptive mechanism consistent with the reduced energetic demands of non-replicating bacteria. Under these conditions, the alternative ETC may be insufficient to fully re-oxidise NADH, thereby inducing malate dehydrogenase activity as a compensatory mechanism. This mechanistic hypothesis would explain the persistence of Mtb under iron-depleted conditions.

Further studies are needed to identify which enzymes of central carbon metabolism are essential for this mechanism. In particular, genetic and biochemical approaches targeting MDH would provide direct evidence for the role of the PCK/PCA-mediated reductive Krebs cycle in malate biosynthesis and secretion under iron-limiting conditions. Additionally, further investigation of the GABA shunt would help clarify its contribution to succinate biosynthesis.

While this study has certain limitations related to the specific experimental conditions used, these observations provide a basis for exploring whether this adaptive mechanism also occurs during infection. An important question is whether metabolites secreted by *Mtb* could substantially modify the surrounding microenvironment, thereby influencing host responses and bacterial pathogenicity.

Mycobacteria are known to exhibit a dynamic interplay between CCM and cell envelope biosynthesis under stress conditions such as hypoxia and antibiotic exposure (Eoh et al., 2022). In this context, the metabolic shift induced by iron starvation may contribute to remodelling of the cell envelope. Consistent with this, iron-starved mycobacterial cells show altered cell envelope thickness and differential expression of genes involved in mycolic acid and cell wall biosynthesis (Vijay et al., 2018; Kurthkoti, 2017). Notably, under iron starvation *Mtb* displays differential sensitivity to antibiotics with distinct structural properties: resistance is observed for smaller molecules such as kanamycin, D-cycloserine, ethionamide, ciprofloxacin, and isoniazid, whereas sensitivity to rifampicin, which has a larger and more complex structure, is maintained (Kurthkoti, 2017). This pattern suggests that cell envelope remodelling under iron starvation may selectively affect antibiotic permeability. Whether such changes also influence direct interactions with host cells, particularly at the bacteria surface-host cell interface remains an important question for future investigation.

## Materials and Methods

### Bacterial strains and growth conditions

In this study H37Rv and Erdman *M. tuberculosis* strains were used. *ΔpckA* and *ΔpckA*::*pckA* (Marrero et al. 2010) (Erdman genomic background) were kindly provided by Professor Sabine Ehrt (Weill Cornell Medicine). The *Δ*icl1 (Lee et al. 2013) strain was kindly provided by Professor Brian VanderVen (Cornell University). The *Δicl1*::*icl1* strain was previously generated (Serafini et al. 2019). For the metabolomic experiments in the Erdman genomic background, the strain *ΔpckA*::*pckA* was used (named Erdman strain in the text and figures and named *ΔpckA*::*pckA* in the comparison with *ΔpckA*). Bacteria were routinely cultivated at 37 °C in 7H9/7H10 medium supplemented with ADC (0.5% bovine serum albumin 0.2% glucose, 0.085% NaCl, 0.003% catalase), 0.05% Tween80 and 0.2% glycerol. Mutant and complemented strains were grown in the presence of selective antibiotics in the 7H9/7H10 pre-cultures. Antibiotics were added at the following concentrations: 20 μg/ml kanamycin (Km); 10 μg/ml gentamicin (Gm); 100 μg/ml hygromycin (Hyg).

For all the experiments bacteria were cultivated in minimal medium (MM; 0.5% w/v KH_2_PO_4_; 0.5 % w/v asparagine; 0.2% glycerol; 0.05% tyloxapol; 0.5% bovine serum albumin 0.2% D-glucose, 0.085% NaCl, pH 6.8) treated with Chelex100 resin (Sigma) at RT for 24 h, replacing Chelex 100 resin after 12 h. The resin was removed by filtration and the medium was supplemented with 40 mg/L MgSO_4_;

0.1 mg/L MnSO_4_ and ZnSO_4_ 0.5 mg/L. For the labelling experiments, 0.2% glycerol was replaced with a mixture of 0.1% ^13^C_3_-glycerol and 0.1% ^13^C_3_-glycerol. The choice of glycerol as the labelled carbon source was based on the superior ability of H37Rv to metabolise this substrate compared to glucose (Serafini et al. 2019).

A source of Fe^3+^ (50 μM of FeCl_3_) was added only in the control condition (High Iron, HI). All the tests of this study were performed in bacterial cells sub-cultured three times (4-7 days of growth) in MM supplemented with (HI) or without (0 μM of FeCl_3_, Low Iron, LI) a Fe^3+^ source. To get growth arrest, twice sub-cultured bacterial cells grown without a source of Fe^3+^ were treated with the siderophore deferoxamine mesylate salt (DFO) at a final concentration of 50 ng/mL^15^. For each experiment parallel HI and LI cultures were also treated. For metabolomics samples, MM was prepared omitting tyloxapol, which interferes with LC-MS analysis (MM No Tyloxapol, MMNT).

All the growth experiments in liquid cultures were performed in standing in T75 (25 mL of culture)/T25 (10 mL of culture) flasks laid down on the incubator floor to optimise the oxygenation, with daily gentle manual shaking.

The input of cells was approximately 1×10^7^ cells/mL for CFU/growth assay and 2×10^7^ cells/mL for all the other analysis (metabolite content, ATP and NADH assays).

Cell viability was assessed by counting Colony Forming Unit (CFU) /mL or by spot density on 7H10/ADC plates. CFU/mL: bacterial culture was serially diluted 10-fold and 50 μL of each dilution was plated in duplicate. Spot’s density: bacterial culture was diluted to 10^7^ cells/mL (OD_600_ ∼0.1) and subjected to four 1:10 serial dilutions (10^6^ to 10^2^ cells/mL); 5 μL of each dilution was spotted on the plate.

### Protein extraction and enzymatic activity

Approximately 10-15 OD_600_ of a bacterial culture were harvested and washed three times with cold PBS 1X and then the pellet resuspended in 600 µL of extraction buffer (DPBS 1X and Complete EDTA-free protease inhibitor cocktail). The cells were disrupted by using acid-washed glass beads (2:1 = buffer volume: beads volume) and a refrigerated beat-beater (6.0 set, 35’’ pulses/1’ pause twice). After centrifugation at 13,000 rpm for 15 min at 4 °C, the supernatant was filtered with a PVDF filter tube before removal from the BLS3 laboratory. Within 6 hrs after extraction, enzymatic activity assays were performed using 10 µl of extract in a 96-well polystyrene microplate, with a final reaction volume of 200 µl. Protein content was determined using the Bradford reagent.

The anaplerotic activity of phosphoenolpyruvate carboxylase was determined as previously described (Machova et al. 2015) coupling the phosphoenolpyruvate carboxylation to oxaloacetate and malate synthesis by using malate dehydrogenase (pig heart, Sigma). The reaction was performed at 37 °C in a buffer containing HEPES/NaOH pH 7.2 100 mM, KHCO_3_ 100 mM, DTT 37 mM, sodium phosphoenolpyruvate 2 mM, GDP 1 mM, MgCl_2_ 2 mM, MnCl_2_ 0.1 mM, malate dehydrogenase 3.5 U/ml, NADH 0.25 mM. The reaction was initiated by adding the MgCl_2_ and MnCl_2_. The decrease of absorbance at 340 nm was measured to monitor NADH oxidation.

Pyruvate carboxylase activity was determined as previously described (Mukhopadhyay & Purwantini 2000) coupling the phosphoenolpyruvate carboxylation to oxaloacetate and malate synthesis by using malate dehydrogenase. The reaction was performed at 37°C in a buffer containing Tris/HCl pH 8.0 50 mM, KHCO_3_ 20 mM, MgCl_2_ 8 mM, di-sodium ATP 8 mM, acetyl-CoA 50 µM, di-sodium NADH 0.2 mM, malate dehydrogenase 3.5 U/ml and sodium pyruvate 20 mM. The reaction was initiated by adding pyruvate. The decrease of absorbance at 340 nm was measured to monitor NADH oxidation.

Malic enzyme activity was determined as previously described (Basu et al. 2018). The reaction was performed at 30 °C in a buffer containing Tris/HCl pH 6.0 100 mM, KHCO_3_ 200 mM, MnCl_2_ 0.5 mM, NAD(P)H 0.6 mM, and sodium pyruvate 50 mM. The reaction was initiated by adding pyruvate. The decrease of absorbance at 340 nm was measured to monitor NAD(P)H oxidation.

Isocitrate lyase activity was determined as previously described (Munoz-Elias et al. 2006). The reaction was performed at 25 °C in a buffer containing MOPS-HCl pH 6.8 50 mM, MgCl_2_ 5 mM, NADH 0.1 mM, 7U L-lactate dehydrogenase (Rabbit). The reaction was pre-incubated 5 minutes at 25 °C and then 1 mM of DL-*threo*-isocitrate was added. The decrease of absorbance at 340 nm was measured to monitor NADH oxidation.

### Metabolite extraction and LC-MS analysis

Cells were sub-cultured twice in MM supplemented with 50 or 0 µM of FeCl_3_, then washed once to remove tyloxapol and resuspended to a final absorbance of 0.2 in 10 mL of MMNT supplemented with 50, 0 µM of FeCl_3_ and 0 µM of FeCl_3_ +DFO. MMNT contained 0.2% of D-glucose, 0.1% of glycerol and 0.1% of (U)-^13^C-labelled glycerol. Cultures were incubated standing at 37 °C. Intracellular polar metabolites were extracted after 8 days by mechanical rupture in an acetonitrile:methanol:water (2:2:1, v/v/v) solution as previously described (Serafini et al. 2019). Extracellular metabolites were extracted after 1, 3 and 8 days by diluting the culture-filtrate 1:5 using cold acetonitrile/methanol (1:1) supplemented with 0.1% formic acid; the samples were vigorously vortexed and stored at -20 °C for 2 hours; they were spun down for 10 minutes at 14,000 rpm and 4 °C, then filtered through 0.22 µm filter tubes.

2 µl of pellet extract and 10 μL of culture-filtrate extract were injected in a 1200 Liquid Chromatography system (Agilent) coupled to an Accurate Mass 6220 TOF (Agilent). Polar elution was performed as previously described using a gradient of two solvents, A (mQ water and 0.1% of formic acid) and B (acetonitrile and 0.1% of formic acid) (Serafini et al. 2019).

The data were analysed by Profinder B.08.00 software and Masshunter Qualitative Analysis B07.00. Intracellular metabolites were normalised by the residual peptide content detected using a BCA assay. Extracellular metabolites were normalised by total ion counts detected by Progenesis software. To plot the data, each normalised value was multiplied or divided for the same number, then the y axis plots report an arbitrary unit.

Due to technical limitations, the phosphorylated intermediates of glycolysis and oxaloacetate could not be analysed.

### ATP and NADH/NAD assays

The ATP assay was performed using the Promega BacTiter-Glo Microbial Cell Viability assay (G8232). Three technical replicates (three aliquots of cells) were performed for each experiment using approximately 10^7^ cells/well (0.1 OD_600_). ATP levels were assessed using an ATP standard curve (μM) and values normalised to OD_600_.The NADH/NAD ratio was determined using the Promega NAD/NADH-Glo assay (G9071). Cells were collected, cooled on ice, and resuspended in a buffer containing PBS, NaOH and dodecyltrimethylammonium bromide according to the kit. The cells were then disrupted with a beat-beater (30’’ 6.5 power at 4 °C) and the extracts were filtered through 0.22 µm filter tubes. The levels of NADH and NAD were then determined according to the kit instructions using NAD and NADH calibration curves. Two technical replicates were performed for each experiment using approximately 3×10^7^ cells. The experiments shown in Figure 1 and Figure 1- figure supplementary 1 were performed in independent experimental sessions, using different batches of reagents and by different operators. These factors account for the differences in the y-axis scale.

### Statistical analysis

Data were expressed as average and standard deviations. Statistical correlations of data were checked for significance using the paired Student T test (p value <0.05).

## Data availability

Metabolomics data used on this study are available via Zenodo (DOI 10.5281/zenodo.18494375). Dataset title: Metabolomics data from *Mycobacterium tuberculosis* partitions the Krebs cycle under iron starvation.

## Acknowledgments

Research funded by H2020 European Commission Program (MSCA-IF GA N. 101029081) and EU funding within the MUR PNRR Extended Partnership initiative on Emerging Infectious Diseases (project no. PE00000007, INF-ACT). We thank Dr Giuseppe Giordano from Pediatric Research Institute Foundation for access to the Progenesis software.

## Author’s contribution

A.S. conceptualised the study, performed experiments, analysed and interpreted data, and wrote the manuscript. A.G. developed the protocol to extract metabolites from culture-filtrate samples, extracted metabolites from the culture-filtrate samples and processed the metabolomics samples in the LC-MS instrument. D.S. performed the isocitrate lyase activity experiment. L.P.S.C. critically reviewed the manuscript. A.G., D.S and R.M. contributed to revise the final draft of the manuscript.

## Supporting Information

### Other supporting materials for this manuscript include the following

Dataset S1

## Extended Results

### Section I. Intracellular and extracellular levels and isotopic distribution analysis of metabolites in LI vs DFO conditions

Few differences were detected between LI and DFO conditions. Unlike DFO, succinate levels do not decrease under LI in both H37Rv and Erdman (Figure 2A **and** Figure 2**–figure supplement 1A**), and (iso)citrate levels do not increase in LI in Erdman strain.

In H37Rv, total labelling analysis shows that labelled succinate levels were similar between LI and HI conditions (Figure 3), while they decreased in DFO, suggesting an alternative route for succinate production under LI. The isotopologue distribution of α-ketoglutarate and succinate in LI show differences compared to DFO in H37Rv (Figure 3). Succinate isotopologue distribution in LI (Figure 3) resemble (iso)citrate, with M+2 more abundant than M+1, whereas α-ketoglutarate as has only M+1 (Figure 3). This suggests that the glyoxylate shunt may be a dominant route only in LI. However, the lack of changes in the intracellular levels (Figure 2) and labelling **(Figure 2 and 3-figure supplement 2D** and data not shown) of (iso)citrate, malate and succinate in the *Δicl1* strain and its complemented strain supports the hypothesis that the glyoxylate shunt is not important under Fe^3+^ starvation. It is possible that in LI compared to DFO: i) α-ketoglutarate may be subject to a different metabolic circuit, and that ii) succinate originates from another route. The GABA shunt may represent this alternative route (Figure 5A **and main text**), which metabolites appear not to be significantly altered under Fe^3+^ starvation (Figure 5B**, E and F**). Like in DFO, M+1 GABA lacks in LI condition (Figure 5C). Overall, the minimal differences observed between LI and DFO conditions suggest that the main metabolic alterations observed in this study are directly attributable to Fe^3+^ starvation, rather than being secondary effects of growth cessation or unspecific DFO effect.

### Section II. Comparison of total ^13^C labelling in intra- and extracellular isocitrate, malate, fumarate and glutamate

A progressive increase in the total ^13^C labelling of extracellular fumarate, glutamate, isocitrate and malate was observed over the course of one week under both DFO and HI conditions (from day 1 to day 8) (Figure 2**-figure supplement 2 A-H**). This temporal rise in ^13^C enrichment likely reflects the gradual incorporation of the labelled carbon through various metabolic pathways, resulting in an expanding pool of labelled metabolites.

When comparing the ^13^C fractions at day 8 between intracellular and extracellular metabolites, in Erdman strain a marked reduction (approximately 50%) of labelling of these metabolites (Figure 2**-figure supplement 2 B, D, F and H,** Figure 4**, and** Figure 5**-figure supplement 2 E**) was observed under DFO conditions. In contrast, in H37Rv, the ^13^C fraction of (iso)citrate and malate remained comparable between intra- and extracellular pools (Figure 3, Figure 2**-figure supplement 2 C and E**), whereas the ^13^C fraction of extracellular glutamate decreased (approximately 50%) like in Erdman strain (Figure 2**-figure supplement 2 G and** Figure 5**-figure supplement 2 B**).

The reduced labelling of glutamate and fumarate is comparable with the metabolic bottleneck downstream of α-ketoglutarate, indicating a slowdown in ^13^C flux beyond this metabolic node. The comparable ^13^C fractions of intra- and extracellular (iso)citrate in H37Rv supports the hypothesis of the block at the α-ketoglutarate level, whereby newly synthesised isocitrate is exported out of the cell rather than being further metabolised through the Krebs cycle. A similar interpretation applies to malate in the H37Rv strain: malate produced from pyruvate and phosphoenolpyruvate (PEP) appears to be secreted into the extracellular space instead of being channelled into downstream pathways. The reduced ¹³C labelling fraction observed in extracellular (iso)citrate and malate, compared to the intracellular pool in the Erdman strain suggests a different regulation of metabolite secretion in this strain. Alternatively, it may be linked to a higher activity of the Krebs cycle and, more broadly, CCM, which could favour the redistribution of newly assimilated carbon throughout the metabolic network. This latter observation may explain the differences in malate isotopologue distribution between H37Rv and Erdman strains.

### Section III. Fumarate synthesis

The ^13^C isotopic profile of fumarate does not align with that of succinate and malate (Figure 4H and L), despite the clear indication that the M+2 malate is derived from succinate (Figure 5H) and that the fumarate route is obligate. The fumarate pool has only the M+1 isotopologue. An explanation for these results may derive from the intracellular localization of the enzymatic complexes involved in the synthesis of malate. Given that succinate dehydrogenase is cell-membrane embedded, it seems reasonable to think that the other enzymes involved in the synthesis and oxidation of succinate are also localised close to the membrane. In this scenario, the succinate-derived fumarate might be immediately transformed to fumarate and difficult to detect. Therefore, the only presence of M+1 isotopologue reveals the activity of an alternative metabolic route whose enzymes are likely localised in a separated area of the bacterial cell. Fumarate is also the degradation product of arginosuccinate via ArgH/Rv1659 lyase (Paul et al. 2013) in the arginine succinate biosynthetic pathway. The entire *Arg* operon is upregulated under Fe^3+^ starvation (Kurthkoti et al. 2017) and our metabolic analysis showed a reduction of M+1 fumarate abundance under Fe^3+^ starvation. The reduction of labelling may represent a symptom of the general reduced cellular metabolism under Fe^3+^ depletion and the upregulation of the pathway a tentative to maintain free fumarate pool size sufficient for cell homeostasis.

Moreover, the presence of only the M+1 fumarate isotopologue, even in the presence of sufficient Fe^3+^, suggests that the specific growth medium used affects the fumarate metabolism. Our previous work (Serafini et al. 2019) demonstrated a clear overlap between the isotopic profiles of fumarate, malate and succinate in cells grown in a media with different composition of C and N sources. The reasons for these differences related to the growth media used remain unclear.

### Section IV. Oxaloacetate

Aspartate is usually used as a proxy for oxaloacetate analysis. However, in our experimental conditions, asparagine (38 mM) is present in the medium, feeding the intracellular aspartate pool. This results in a consistent amount of unlabelled Asn-derived oxaloacetate entering the Krebs cycle, and therefore aspartate cannot be used to track carbon flux. The reduced labelling percentage of aspartate (below 5%, data not shown) compared to the other Krebs cycle intermediates confirms that. The consistent amount of unlabelled Asn-derived oxaloacetate explains the prevalence of M+1 and M+2 isotopologues in (iso)citrate, α-ketoglutarate and succinate (Figure 3 and 4) which is likely due to PDH activity (**Figure 3 and 4-figure supplement 1A-C**).

### Section V. Comparison of theoretical scenarios from Figure 3 and 4-figure supplement 1D-S with labelling results

Only the scenarios illustrated in the **Figure 3 and 4-figure supplement 1D, E, P and Q** lead to the production of M+3 malate. Despite producing M+3 malate, the **Figure 3 and 4-figure supplement 1D** scenario also produces M+5 isocitrate and M+4 succinate, which show low abundance in our measurements (Figure 3 **and 4**), suggesting that additional route/s contributes to the generation of M+3 malate. Importantly, this scenario employs two labelled acetyl-CoA molecules. Since the abundance of labelled pyruvate, the precursor of labelled acetyl-CoA, is well below 50% of the total pyruvate (Figure 3 **and 4**), this scenario is unlikely.

The **Figure 3 and 4-figure supplement 1E** scenario produces M+3 malate, M+2 succinate and M+3 (iso)citrate. M+2 succinate is significantly abundant in HI and DFO (Figure 3 **and 4**), whereas M+3 (iso)citrate increases in DFO condition (Figure 3 **and 4**) suggesting that this last isotopologue can derive from PCK-derived oxaloacetate. The **Figure 3 and 4-figure supplement 1E** scenario employs one labelled acetyl-CoA that is used by malate synthase (GlcB), similarly to Figure 3 **and 4–figure supplement 1F** scenario that employs one labelled acetyl-CoA but that is used by citrate synthase (GltA). The probability of the **Figure 3 and 4-figure supplement 1E** scenario should be equivalent to that of the **Figure 3 and 4-figure supplement 1F** scenario. The latter produces M+1 malate, which abundance decreases in DFO, M+5 (iso)citrate and M+4 succinate, which are the least abundant isotopologues of (iso)citrate and succinate and, their abundance slightly increases in DFO (**Figure. 3 and 4**). Additionally, the isotopologues produced in the **Figure 3 and 4-figure supplement 1F** combination in DFO are less abundant in our experimental data compared to those produced in the **Figure 3 and 4-figure supplement 1E** combination.

The Figure 3 **and 4–figure supplement 1P** scenario yields M+3 malate, M+2 succinate and M+3 (iso)citrate, however it derives from the combination of two labelled acetyl-CoA, making it as unlikely as the Figure 3 **and 4–figure supplement 1D** scenario.

Finally, the **Figure 3 and 4-figure supplement 1Q** scenario produces M+1 (iso)citrate, together with M+3 malate. This scenario includes one labelled acetyl-CoA that is used by GlcB, similarly to **Figure 3 and 4-figure supplement 1R** scenario that employs one labelled acetyl-CoA but that is used by citrate synthase (GltA). The probability of the **Figure 3 and 4-figure supplement 1Q** scenario should be equivalent to that of the **Figure 3 and 4-figure supplement 1R** scenario. The latter produces M+1 malate, M+2 succinate and M+3 (iso)citrate. Both **Figure 3 and 4-figure supplement 1Q** and **Figure 3 and 4-figure supplement 1R** produce specific isotopologues whose changes in abundance in the DFO condition compared to the HI condition show opposite directions (Figure 3 **and 4**).

In summary, while the four theoretical scenarios that include the glyoxylate cycle described above reflect qualitatively our labelling results, several quantitative discrepancies exist with the measured and predicted abundances of the isotopologues. These observations do not explain the increased abundance of M+3 malate consistently observed under Fe^3+^ starvation and raise doubts about the relevance of glyoxylate shunt under Fe^3+^ starvation.

## Supplementary Figures

**Figure 1 – figure supplement 1.**
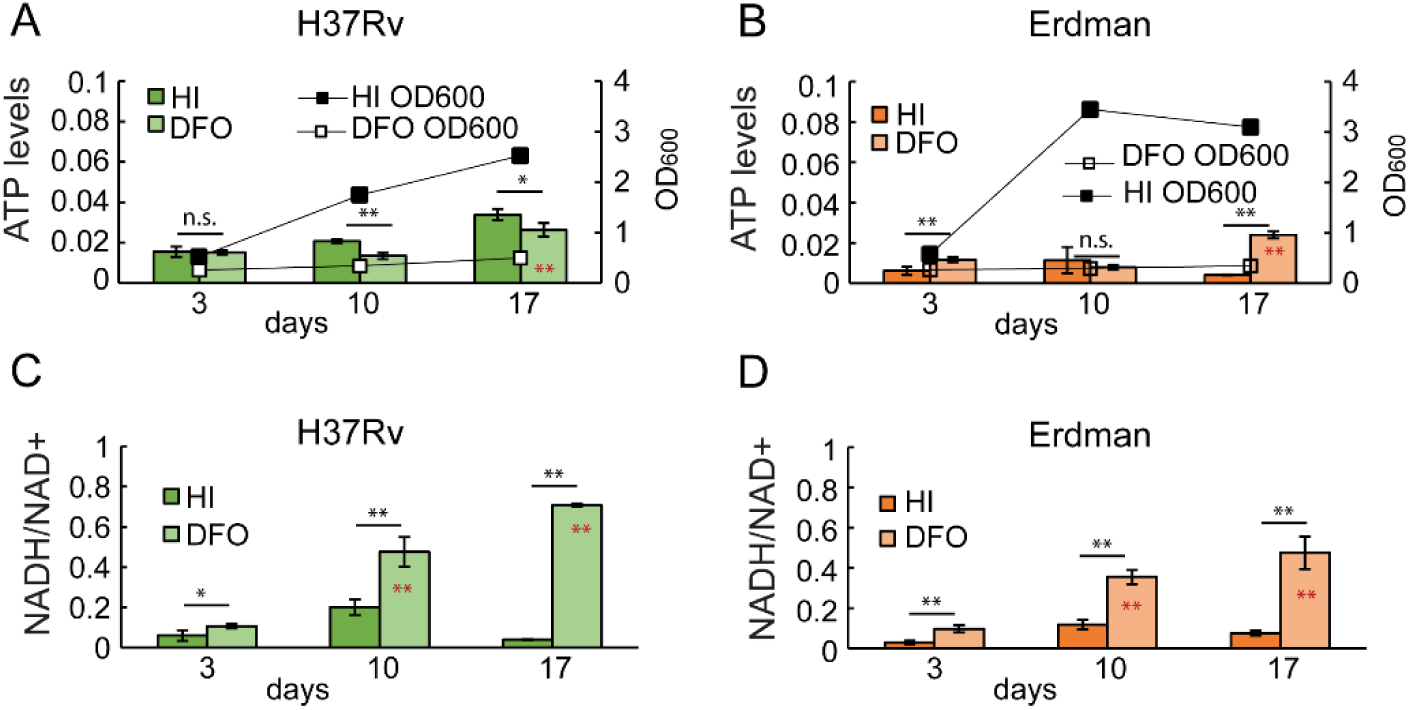
Intracellular ATP content and NADH/NAD+ ratio in H37Rv and Erdman strain. Cells were exposed to 50 μM FeCl_3_ (HI: High Iron) or 0 μM FeCl_3_+ DFO (DFO). Samples were collected after 3, 10 and 17 days of exposure. A, B) The charts show ATP levels and growth (OD_600_). ATP levels were calculated as µM of ATP molecules in 10^7^ cells (0.1 optical density at 600 nM). C, D) The charts show the ratio between NADH and NAD levels. NADH and NAD+ levels were normalised on protein content of the extract. All the data are the average and standard deviation of two biological replicates, two culture aliquots from each replicate and two technical replicates on the 96-well plate. N.s. = non-significant, p value > 0.05. * = p value < 0.05. ** = p value < 0.01. Black asterisk (outside the bar): DFO vs HI; red asterisk (inside the bar): day 17 vs day 10 and day 3, and day 10 vs day 3.

**Figure 2 - figure supplement 1.**
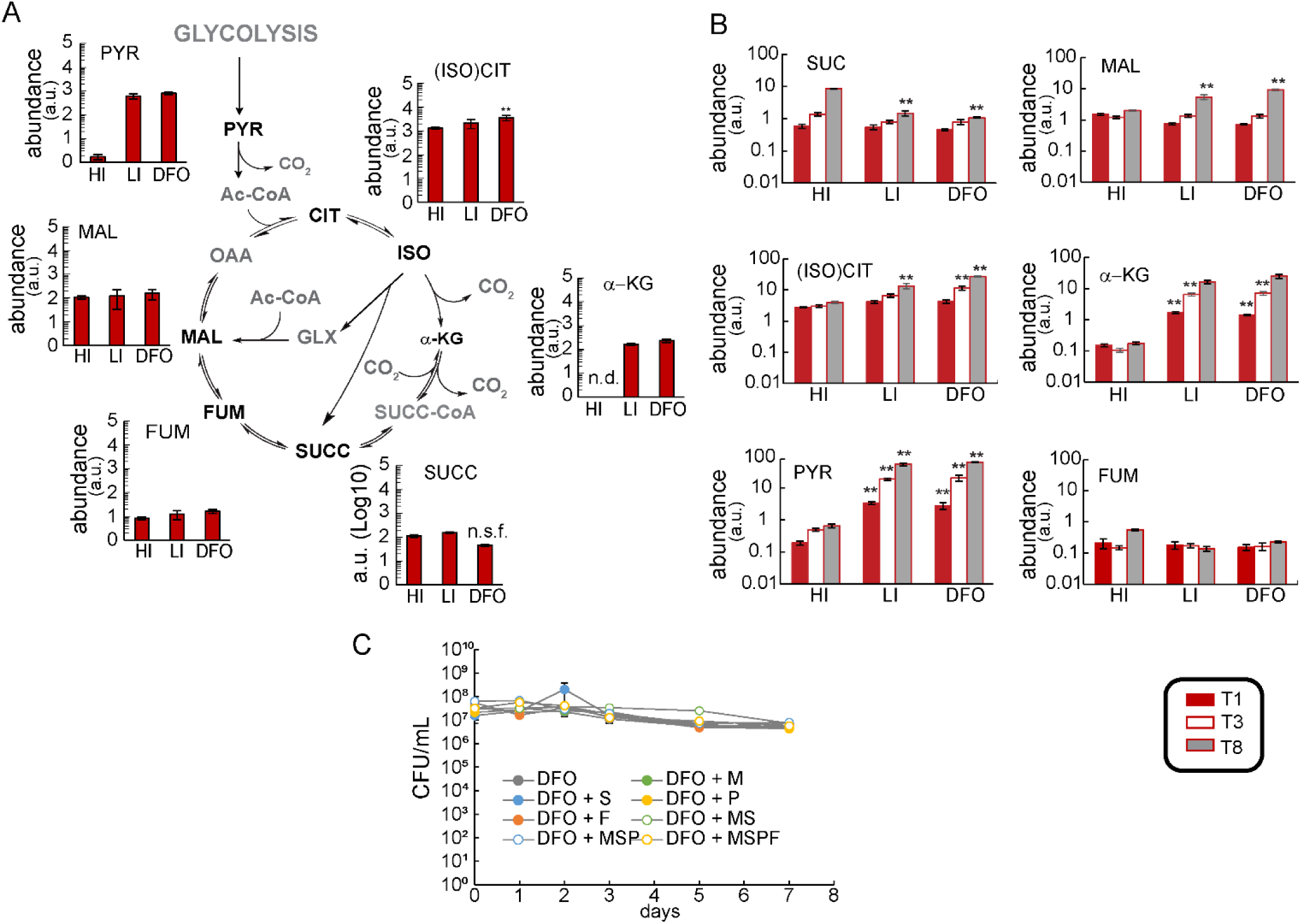
Intracellular and extracellular levels of metabolites in the Erdman strain. Cells were exposed to 50 μM FeCl_3_ (HI: High Iron), 0 μM FeCl_3_ (LI: Low Iron,) or 0 μM FeCl_3_+ DFO (DFO). The analysed metabolites are shown in black in the schematic pathways. A) Intracellular polar metabolites levels at 8 days. the y-axis is shown on Log10 scale tick labels (0–5) represent the exponents of 10 (10⁰–10⁵); values are reported in arbitrary units. The slight decrease in succinate levels (0.66 FC) was not observed in the other four independent experiments in which no changes were observed in LI and DFO conditions compared to HI condition. B) Extracellular polar metabolites levels at 1,3 and 8 days. The plots show the normalised levels of metabolites. The y-axis is shown on is in Log_10_ scale, values are reported in arbitrary units. The data represent the average and the standard deviation of four biological replicates from an independent experiment, representative of four independent experiments for the HI and DFO conditions, and two independent experiments for the LI condition. The y axis is in Log_10_ scale, an arbitrary unit is reported. G) Viability of cells exposed to DFO in the presence of 2 mM of malate (DFO+M), succinate (DFO+S), pyruvate (DFO+P) or a combination of the three (DFO+M+S+P). The plot reports the viability measured as CFU/mL in Log_10_ scale. The data are representative of one independent experiment (average and average deviation of two technical replicates). The p-values were calculated against the HI condition and independently for the four experiments; the highest p-value is reported. * = p value < 0.05; ** = p value <0.01. n.d. = not detected. n.s.f. = not significant fold change, the observed trend change was different between independent experiments. Ac-CoA: acetyl-CoA; CIT: citrate; FUM: fumarate; α-KG: α-ketoglutarate; ISO: isocitrate; (ISO)CIT: isocitrate and citrate; MAL: malate; OAA: oxaloacetate; PYR: pyruvate SUCC: succinate; SUCC-CoA: succinyl-CoA.

**Figure 2 - figure supplement 2.**
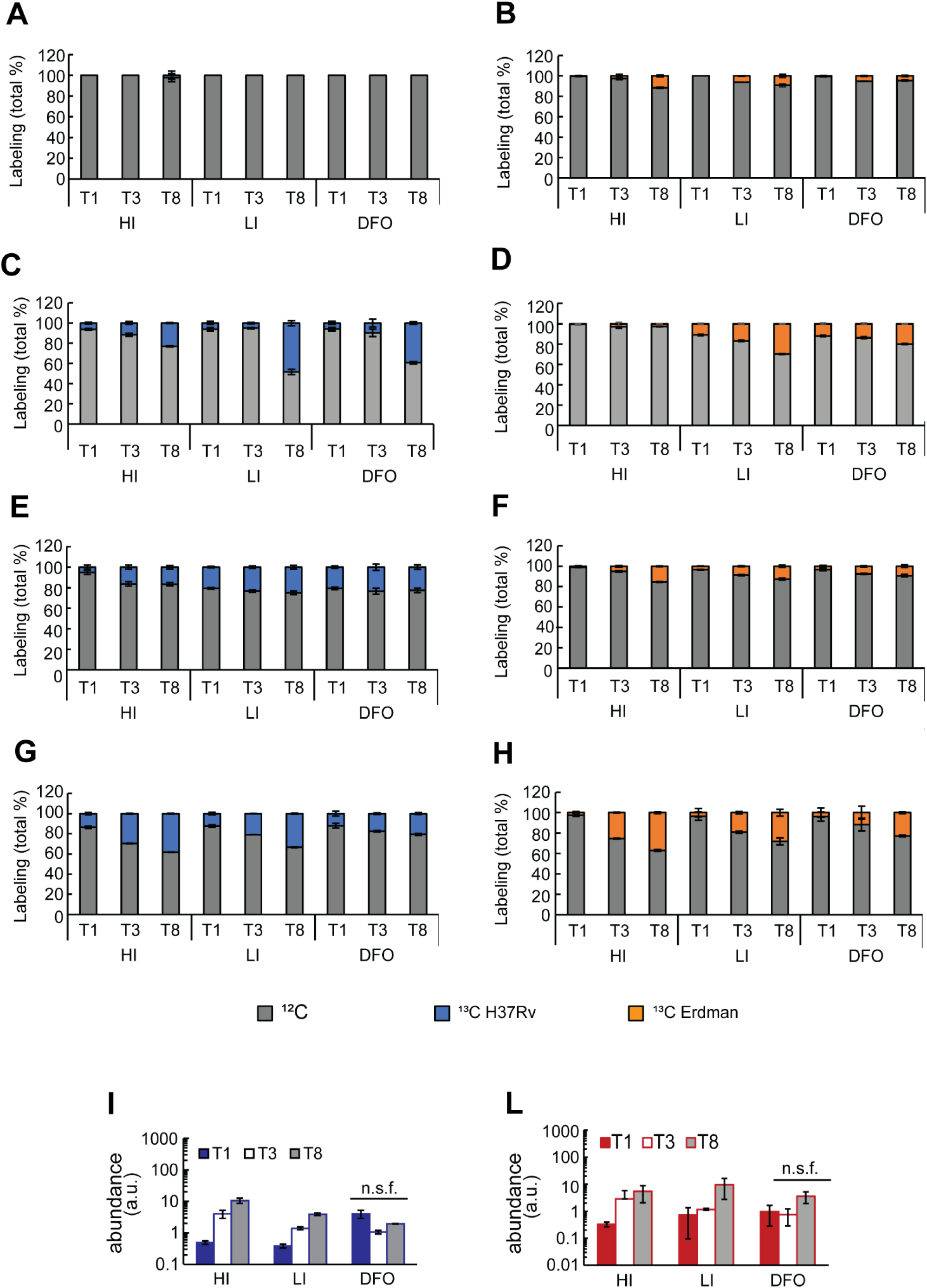
Abundance and ^13^C labelling of extracellular metabolites. Cells were exposed to 50 μM FeCl_3_ (HI: High Iron), 0 μM FeCl_3_ (LI: Low Iron,) or 0 μM FeCl_3_+ DFO (DFO). Metabolites were extracted from culture-filtrate at 1, 3 and 8 days in H37Rv (A,C,E,G,I) and Erdman strain (B,D,F,H,L). A-F) ^13^C total labelling of fumarate (A,B), (iso)citrate (C,D) malate (E,F) and glutamate (G,H). I,L). The plots show the normalised levels of glutamate; the y axes is in log_10_ scale. The data represent the average and the standard deviation of four biological replicates from an independent experiment, representative of two independent experiments. n.s.f. = not significant fold change, the observed trend change was different between independent experiments; a.u. = arbitrary unit.

**Figure 2 - figure supplement 3.**
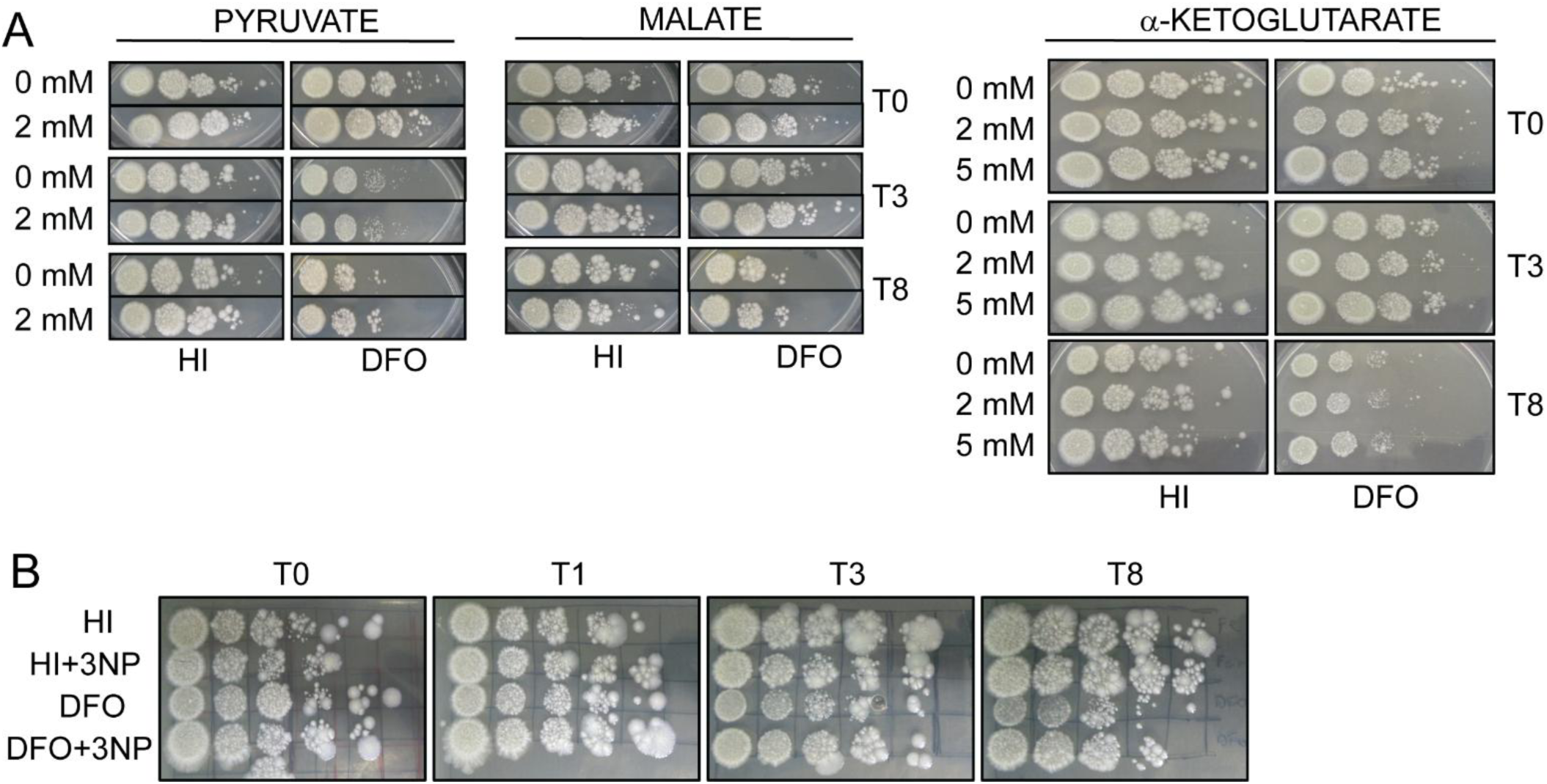
Viability of *Mtb* exposed to severe Fe^3+^ starvation and in the presence of 2 mM or 5 mM Krebs cycle intermediates (A, H37Rv) or 200 μM 3NP (B, Erdman). Cells were exposed to 50 μM FeCl_3_ (HI: High Iron), or 0 μM FeCl_3_+ DFO (DFO) in liquid medium for 8 days. Aliquots of cells were collected after 0, 1, 3 and 8 days, diluted to a final OD_600_ of 0.1 and 5 μL of a 10-fold serial dilution were plated on 7H10/ADC. Growth was recorded after 19-25 days.

**Figure 2 and 3 - figure supplement 1.**
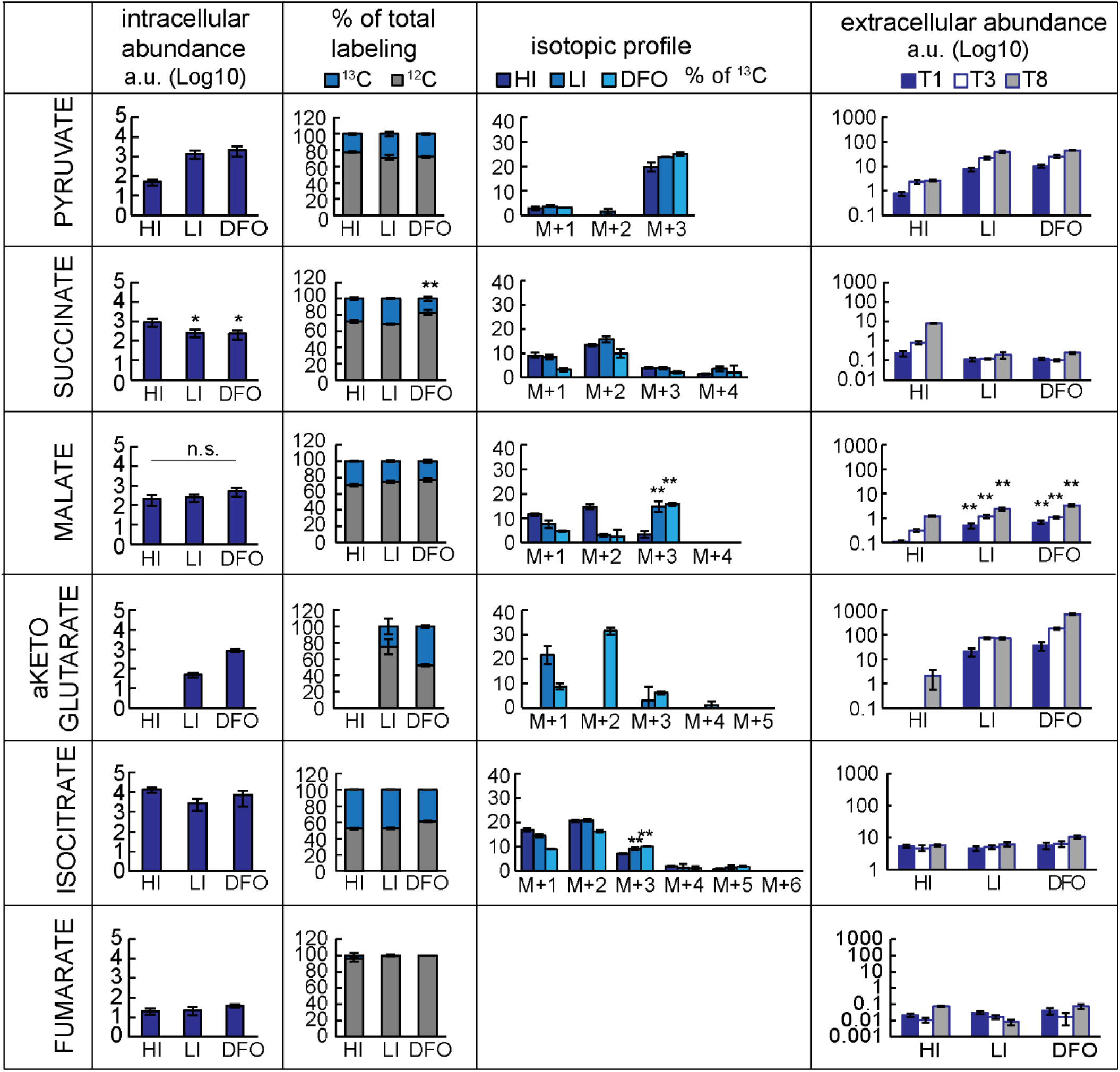
Analysis of metabolites in the H37Rv-derived *Δicl1::icl1* complemented strain. Cells were exposed to 50 μM FeCl_3_ (HI: High Iron), 0 μM FeCl_3_ (LI: Low Iron,) or 0 μM FeCl_3_+ DFO (DFO), and fed with ^13^C_3_-glycerol. Intracellular metabolites were determined after 8 days; extracellular metabolites were determined after 1,3 and 8 days. The plots show the normalised levels of metabolites. The data represent average and standard deviation from one experiment and four biological replicates. For intracellular abundance plots, the y-axis is shown on Log_10_ scale tick labels (0–5) represent the exponents of 10 (10⁰–10⁵), values are reported in arbitrary units. For extracellular abundance plots. The y-axis is shown on is in Log_10_ scale, values are reported in arbitrary units. The column headers contain the labels of the y axis. The p-values were calculated against HI condition. n.s. = non-significant, p value >0.05; * = p value <0.05; ** = p value <0.01. a.u. = arbitrary unit.

**Figure 2 and 3 - figure supplement 2.**
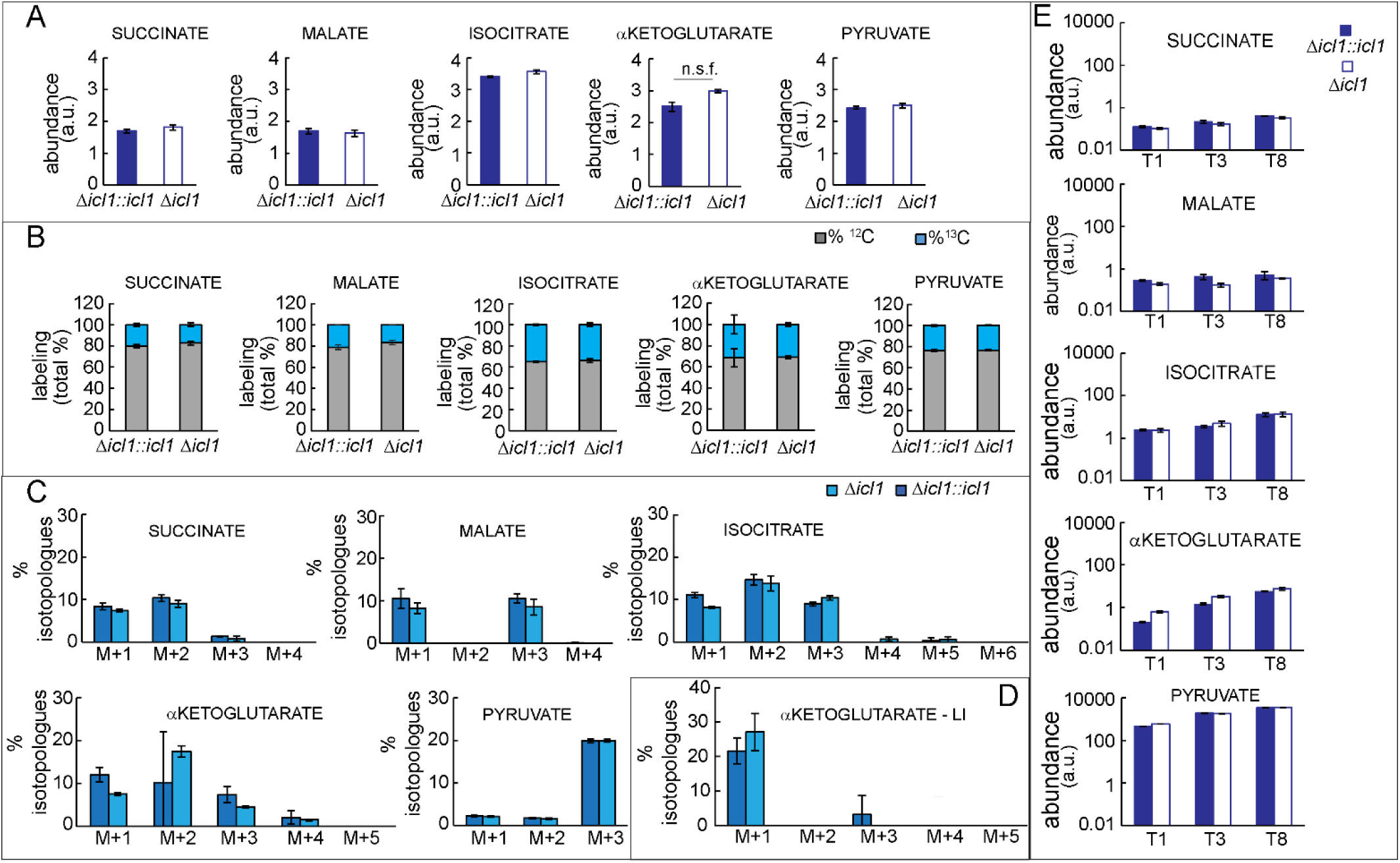
Analysis of metabolites in the H37Rv-derived *icl1* mutant *(Δicl1*) compared to its complemented strain (*Δicl1::icl1*) in the DFO condition. Cells were exposed to 0 μM FeCl_3_+ DFO (DFO), and fed with ^13^C_3_-glycerol for 8 days. Intracellular metabolites were extracted after 8 days; extracellular metabolites were extracted after 1,3 and 8 days. A) Normalised abundance of intracellular metabolites, the y-axis is shown on Log_10_ scale tick labels (0–4) represent the exponents of 10 (10⁰–10^4^), values are reported in arbitrary units. B) Total percentage of labelled and unlabelled metabolites. C) Isotopologue distribution of metabolites. D) Isotopologue distribution of the α-ketoglutarate pool in the low iron (LI; 0 μM FeCl_3_) condition. E) Normalised abundance of extracellular metabolites, the y-axis is shown on is in Log_10_ scale, values are reported in arbitrary units. A, B, C, E) Data from one experiment representative of two independent experiments and four biological replicates. D: data from one independent experiment and four biological replicates each. The data represent average and standard deviation of four biological replicates from an independent experiment. n.s.f. = not significant fold change, the observed trend change was different between independent experiments. a.u. = arbitrary unit.

**Figure 3 and 4 - figure supplement 1.**
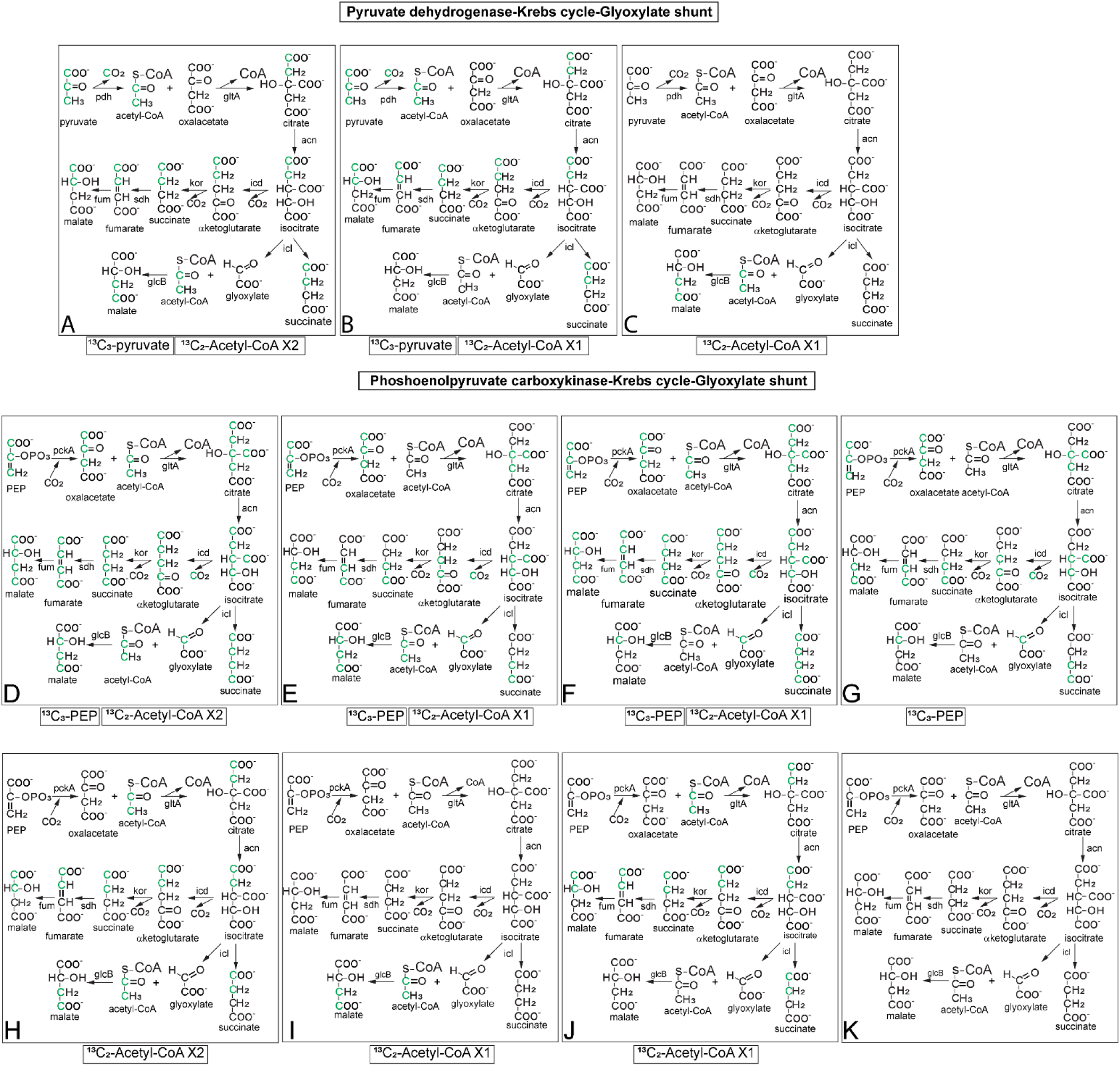
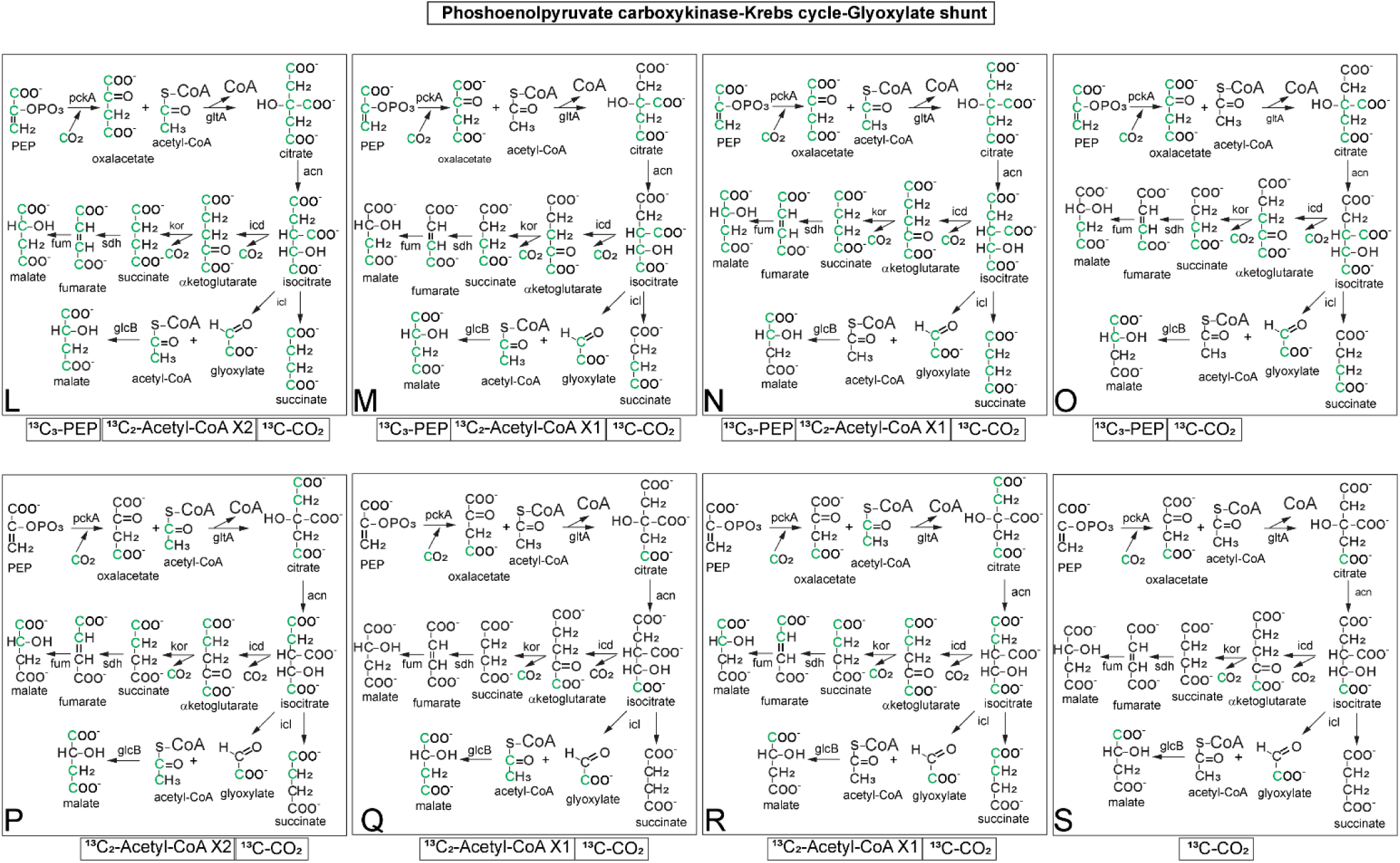
Schematic representation of ^13^C-phosphoenolpyruvate, ^13^C-pyruvate and ^13^C-carbon dioxide assimilation through PDH, PCK, the Krebs cycle and the glyoxylate shunt. Atoms of ^13^C are marked in green. The panels A-C include the activity of pyruvate dehydrogenase (PDH), the Krebs cycle and the glyoxylate shunt. The panels D-S include the activity of phosphoenolpyruvate carboxykinase (PCK), the Krebs cycle and the glyoxylate shunt. Labelled acetyl-CoA is assumed to derive from PDH activity. The bottom of each panel indicates which input metabolite is labelled. The labelling profiles of metabolites downstream to phosphoenolpyruvate (PEP) are identical if PCK is replaced by pyruvate carboxylase and PEP is replace by pyruvate. To simplify, only one of the enzymes for α-ketoglutarate degradation and citrate synthesis are reported. Acn: aconitase. GlcB: malate synthase. GltA: citrate synthase. Icd: isocitrate dehydrogenase. Icl: isocitrate lyase. Kor: αketoglutarate ferredoxin-oxidoreductase. Mdh: malate dehydrogenase. Sdh: succinate dehydrogenase.

**Figure 3 and 4 - figure supplement 2.**
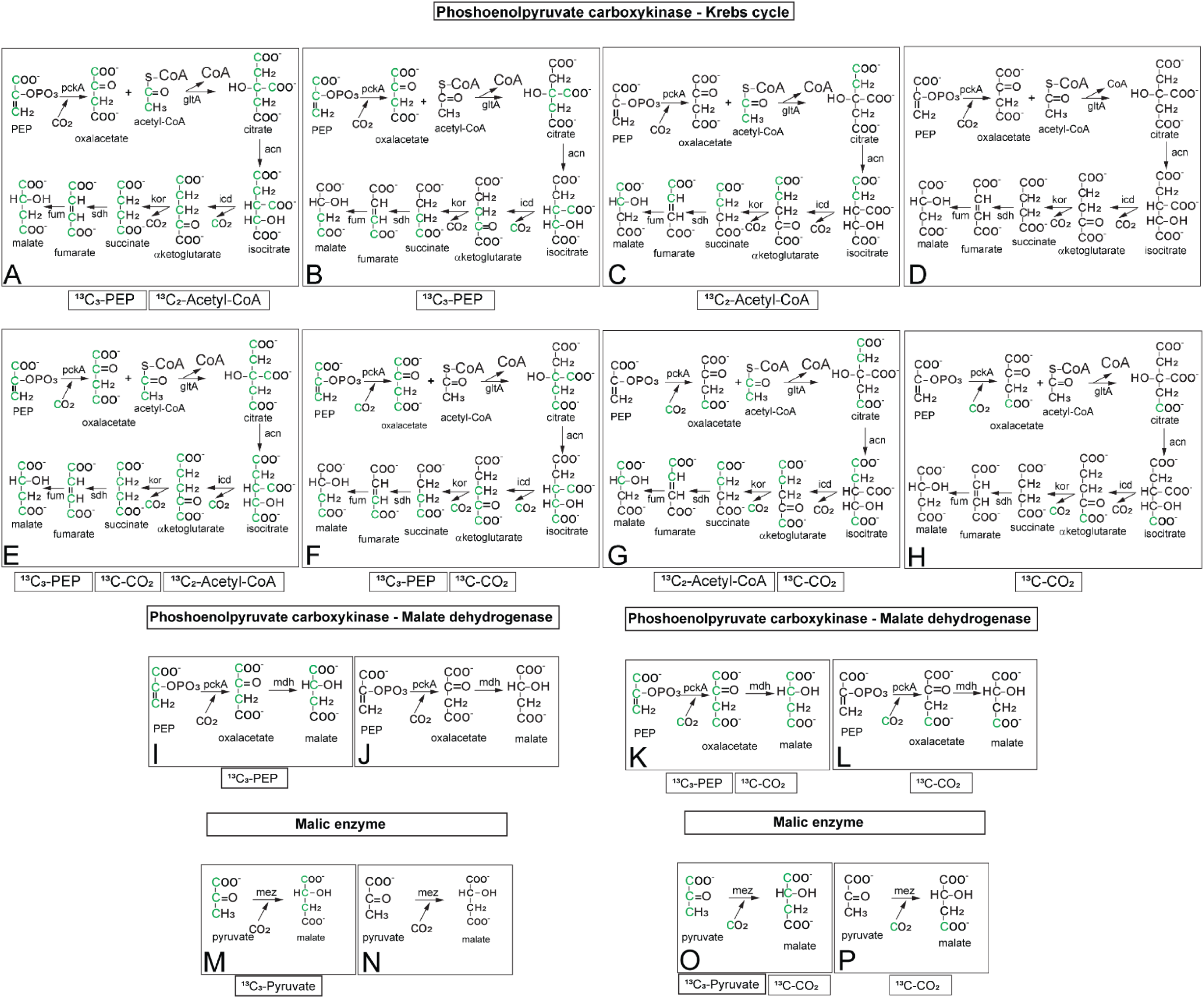
Schematic representation of ^13^C-phosphoenolpyruvate, ^13^C-pyruvate and ^13^C-carbon dioxide assimilation through PCK, MEZ and the Krebs cycle. The panels A-H include the activity of phosphoenolpyruvate carboxykinase (PCK) and the Krebs cycle. Labelled acetyl-CoA is assumed to derive from PDH activity. The panels I and L include the anaplerotic reaction of PCK and the reduction of oxaloacetate to malate by MDH. The labelling profiles of metabolites downstream to phosphoenolpyruvate (PEP) are identical if PCK is replaced by pyruvate carboxylase and PEP is replaced by pyruvate (A-L). The panels M-N include the anaplerotic reaction of malic enzyme (MEZ) from pyruvate to malate. To simplify, only one of the enzymes for α-ketoglutarate degradation and citrate synthesis are reported. The bottom of each panel indicates which input metabolite is labelled. Acn: aconitase. GltA: citrate synthase. Icd: isocitrate dehydrogenase. Kor: α-ketoglutarate oxidoreductase. Sdh: succinate dehydrogenase.

**Figure 3 and 4 - figure supplement 3.**
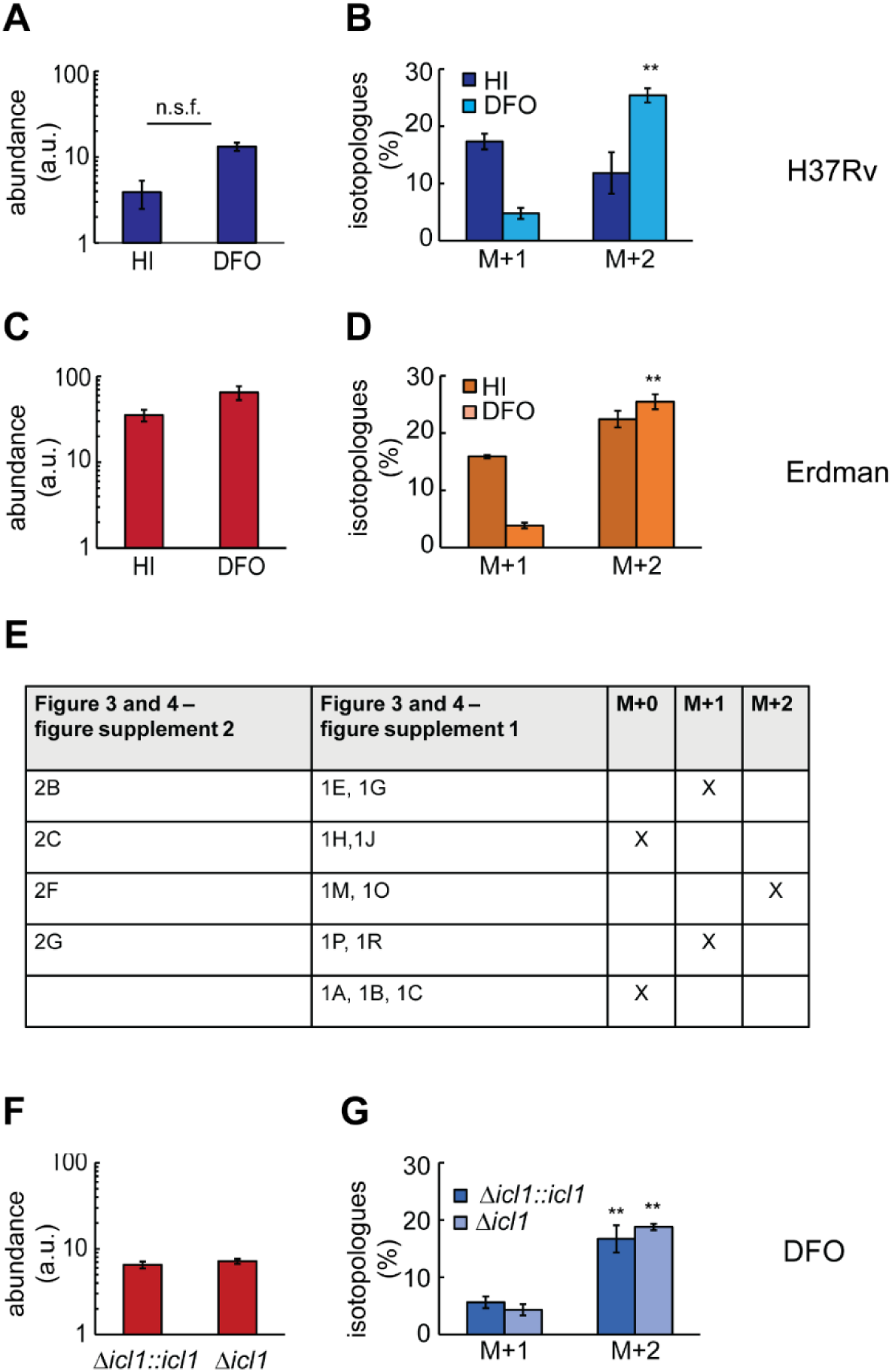
Analysis of intracellular glycine in the H37Rv, Erdman and H37Rv-derived icl1 mutant in HI and DFO condition. Cells were exposed to 50 μM FeCl_3_ (HI: High Iron), 0 μM FeCl_3_+ DFO (DFO), and fed with ^13^C_3_-glycerol for 8 days. A, C) Normalised levels of glycine in H37Rv and Erdman strains; the y-axis is shown on Log_10_ scale tick labels (0–5) represent the exponents of 10 (10⁰–10⁵), values are reported in arbitrary units. B, D) Isotopologue distribution expressed in percentage in H37Rv and Erdman strains. Plots show average and standard deviation from one experiment and four biological replicates. E) The table shows the qualitative analysis of the glycine isotopologues produced by the scenarios illustrated in the figure 3 and 4 – figure supplement 1 and 2. F) Normalised levels of glycine in an icl1 mutant and complemented strains; the y-axis is shown on is in Log_10_ scale, values are reported in arbitrary units. G) Isotopologue distribution expressed in percentage in an *icl1* mutant and complemented strains. The data are representative of two-four independent experiments. The p-values were calculated against HI condition. a.u.: arbitrary unit (see methods paragraph). n.s.f = non-significant fold change, the observed trend change was different between independent experiments. ** = p value <0.01.

**Figure 5 - figure supplement 1.**
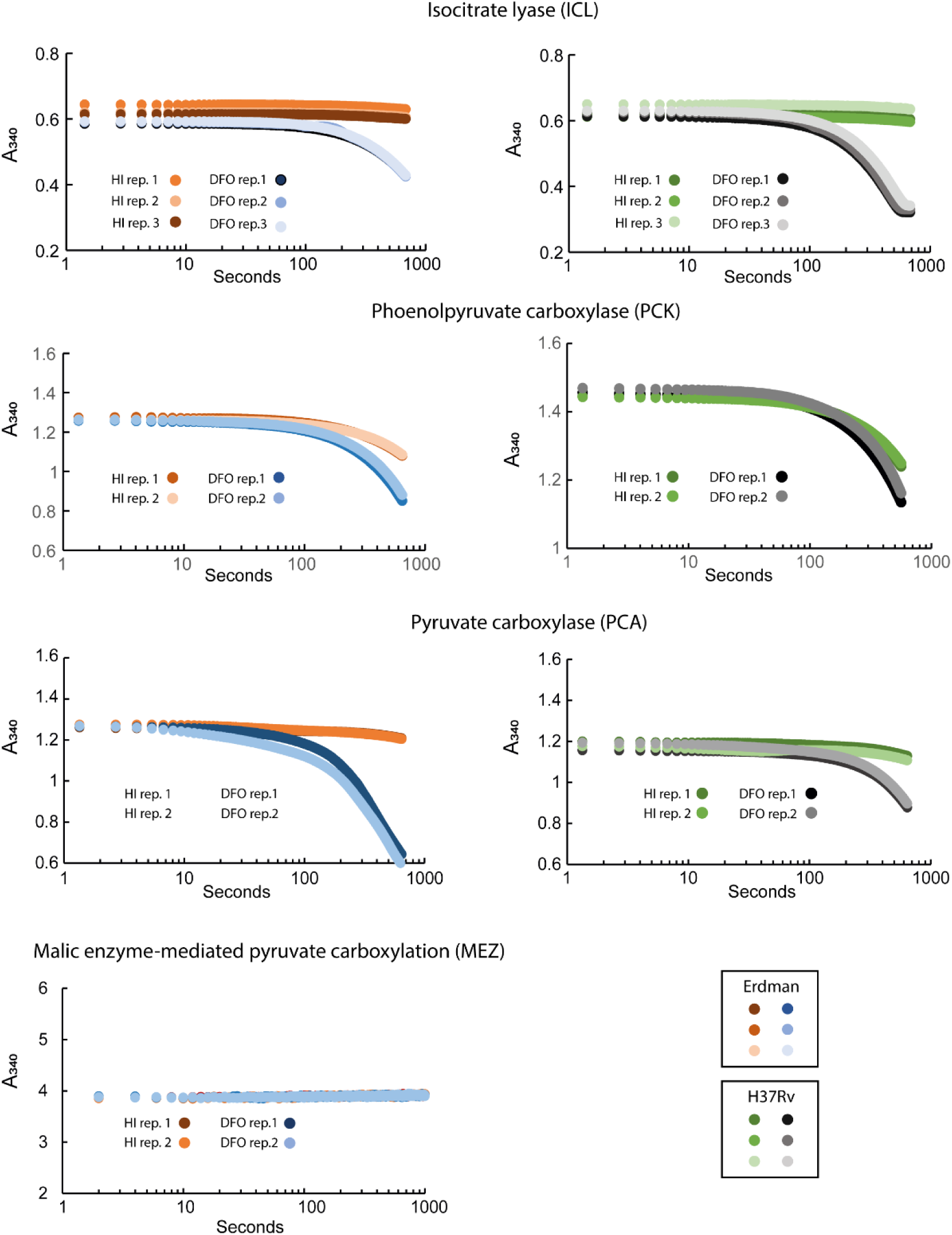
ICL, PCK, PCA and MEZ activities in H37Rv and Erdman strains. The plots report the decrease of absorbance at 340 nm due to the use of NAD(P)H or NADH from two or three technical replicates. The data are representative of one (ICL, MEZ) or two (PCK, PCA) independent experiments. The activity was performed in protein extracts from cells exposed to 50 μM FeCl_3_ (HI: High Iron), and 0 μM FeCl_3_+ DFO (DFO) for 3 days. The plots report the raw values not normalised on protein content.

**Figure 5 - figure supplement 2.**
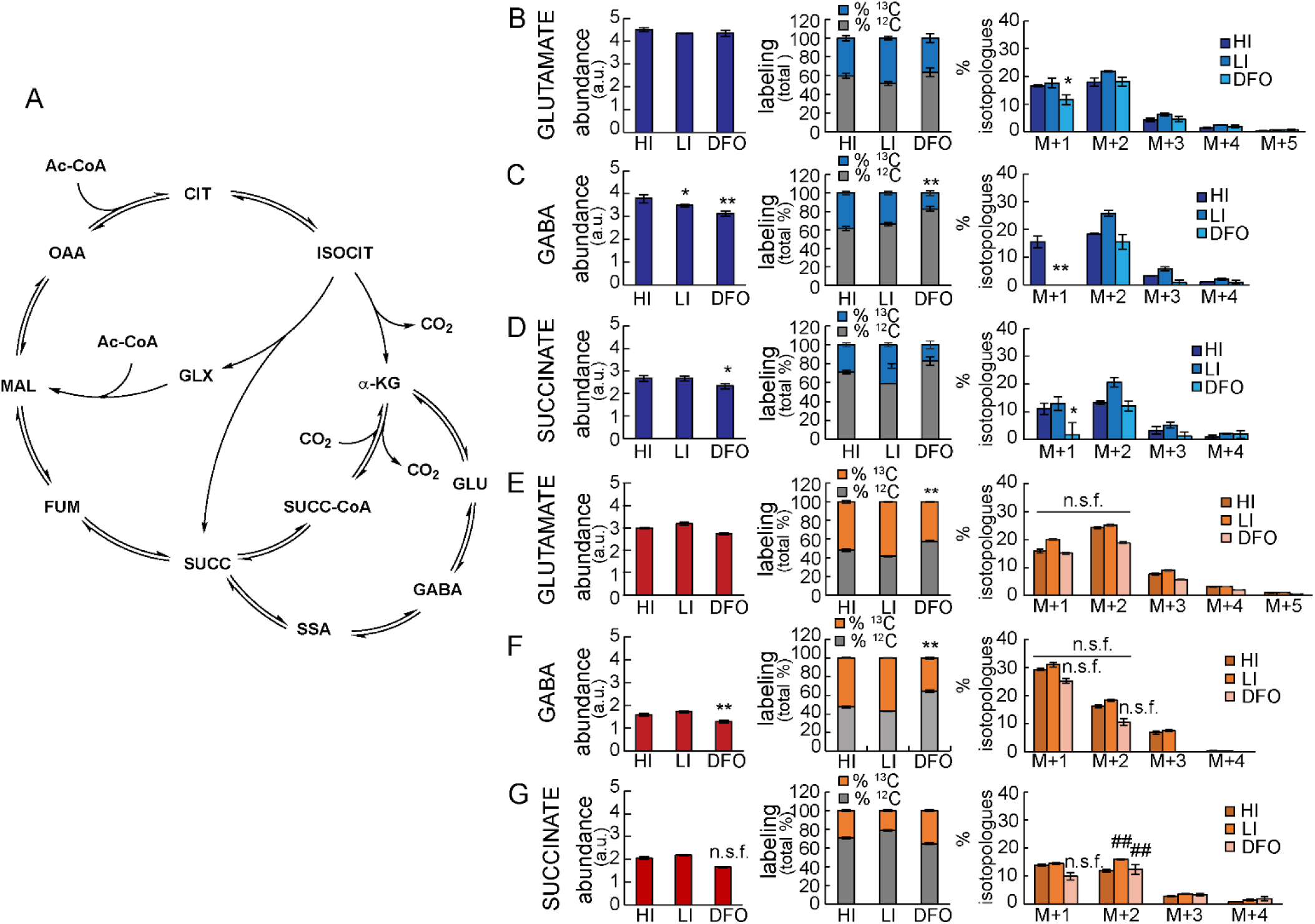
Analysis of GABA shunt metabolites in H37RV and Erdman strains. Cells were exposed to 50 μM FeCl_3_ (HI: High Iron), 0 μM FeCl_3_ (LI: Low Iron,) or 0 μM FeCl_3_+ DFO (DFO), and fed with ^13^C_3_-glycerol for 8 days. A) Schematic representation of the GABA shunt and the Krebs cycle. B, C, D) H37Rv; E, F, G) Erdman strain. B, E) glutamate; C, F) gamma-aminobutyric acid (GABA); D, G) succinate. Plots with one color-filled histograms represent the normalised levels of metabolites the y-axis is shown on Log_10_ scale tick labels (0–5) represent the exponents of 10 (10⁰–10⁵), values are reported in arbitrary units; plots with two color-stacked histograms represent the total abundance in percentage of labelled and unlabelled metabolite pools; plots with clustered histograms represent the abundance in percentage of each isotopologue. H37Rv data are from one experiment representative of two independent experiments and four biological replicates each; Erdman-derived strain data are from one experiment representative of four independent experiments (only two of them for the LI condition) and four biological replicates each. The histograms show average and standard deviation of four biological replicates from an independent experiment. The p-values were calculated against the HI condition; the highest p-value from the two independent experiments is shown. The differences between M+1 and M+2 isotopologues in the Erdman strain (E, F) are not significant between the three experiments, the trend varied between independent experiments. a.u.: arbitrary unit (see methods paragraph). n.s.f.= non-significant fold change, the observed trend change was different between independent experiments. DFO vs HI or LI vs HI: * =p value <0.05; ** =p value <0.01; “n.s.f.” above the individual M+1 and M+2 bars of succinate and GABA.

